# Glycomimetic Lysosome-targeting Chimeras (GLYTACs) for Degradation of Growth Factors and Growth Factor Receptors in Cancer Cells

**DOI:** 10.64898/2025.12.30.696977

**Authors:** Julianna L. Follmar, Toshiki Tabuchi, Kamil Godula

## Abstract

Many cancers depend on extracellular growth factors within the tumor microenvironment to drive aberrant signaling, proliferation, and survival. For example, anticancer therapies targeting vascular endothelial growth factor activity have been effective in blocking pro-angiogenic and pro-growth signals, including receptor tyrosine kinase inhibitors (e.g., Sunitinib and Sorafenib) and monoclonal antibodies (e.g., Bevacizumab). However, the effectiveness of these therapies is frequently limited by compensatory growth factor signaling and incomplete blockade, contributing to drug resistance and suboptimal long-term responses. To address these challenges, we developed Glycomimetic Lysosome-targeting Chimeras (GLYTACs) that sequester and degrade extracellular growth factors in the cancer cell environment. GLYTACs exploit growth factor interactions with cell-surface heparan sulfate (HS) glycans, a feature shared by many pro-tumorigenic signals and their receptors, by combining an HS-glycomimetic arm for growth factor binding with a polyvalent glycopolymer ligand targeting the lysosomal recycling cation-independent mannose 6-phosphate receptor (CI-M6PR) for efficient internalization and degradation. Treating HeLa cells with a heparin-based GLYTAC prototype drove rapid uptake and degradation of extracellular fibroblast growth factor 2 (FGF2). The capacity of heparin to promote the association of FGF2 with its cognate receptors (FGFRs) led to the degradation of the entire receptor-ligand complex, thereby reducing the availability of FGFRs at the cancer cell surface, which are necessary for sustained pro-oncogenic signaling. These findings highlight the potential of GLYTACs as an alternative to existing growth factor-blocking anticancer therapies and as a strategy to reshape the extracellular signaling environment of tumors.

## INTRODUCTION

The tumor microenvironment is rich in extracellular biochemical cues that regulate critical processes, including cell proliferation, survival, and angiogenesis.^1^ For instance, fibroblast growth factors (FGFs)^2^ and vascular endothelial growth factor (VEGF)^3^ drive malignant progression by sustaining tumor cell proliferation and vascularization. Disruption of these pathways has emerged as an important anticancer therapeutic strategy.^4,5^ Examples of promising therapeutics include the receptor tyrosine kinase inhibitors Sunitinib and Sorafenib^6^, and the monoclonal antibody Bevacizumab,^7^ which neutralize VEGF and its associated signaling pathways, thereby suppressing tumor expansion and vascularization. However, the clinical impact of these therapies is frequently undermined by compensatory signaling,^8^ incomplete target blockade,^9^ and the tumor niche’s dynamic adaptability.^10^ As a result, drug resistance and suboptimal inhibition of oncogenic pathways remain persistent challenges.^11,12^

We propose an alternative strategy to current receptor-blocking therapies that limits pro-oncogenic signaling by removing extracellular growth factors from the tumor microenvironment. This strategy exploits a shared feature of many prooncogenic factors, such as FGFs and VEGFs: their reliance on cell-surface heparan sulfate (HS) glycan binding,^13,14,15^ which can be harnessed to promote their lysosomal degradation and clearance (**Figure 1**).

**Figure 1.**
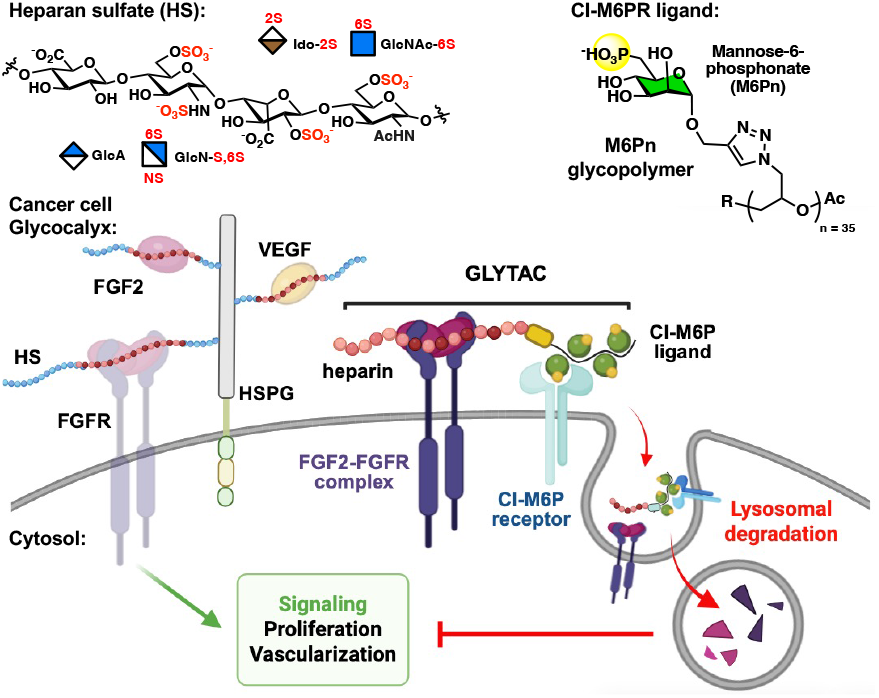
Glycomimetic Lysosome-Targeting Chimeras (GLYTACs) as degraders of pro-oncogenic extracellular signals. GLYTACs exploit interactions between pro-growth and pro-angiogenic signals (e.g., FGFs or VEGFs) and their receptors, and with cell-surface heparan sulfate (HS) glycans, to drive lysosomal uptake, degradation, and clearance within the cancer cell microenvironment. The GLYTAC design incorporates a heparin glycodomain to capture signaling proteins and a glycopolymer ligand for the lysosomal-recycling cation-independent mannose-6-phosphate receptor (CI-M6PR), thereby promoting internalization and degradation.

Inspired by lysosome-targeting chimeras (LYTACs) introduced by Banik and Bertozzi for the targeted degradation of cell surface proteins,^16^ which employ antibodies fused to a ligand for the lysosome-recycling calcium-independent mannose-6-phosphate receptor (CI-M6PR), we aimed to develop a new class of glycomimetic LYTACs, called GLYTACs, which combine a CI-M6PR ligand with an HS glycan-mimetic arm to promote the capture and degradation of extracellular growth factors (**Figure 1**).

Here, we introduce a heparin-based GLYTAC prototype and summarize experimental results demonstrating its ability to sequester and promote lysosomal degradation of FGF2 and its receptor, FGFR1, in HeLa cells. Many extracellular signaling proteins beyond FGF2 interact with cellular HS, providing an opportunity to simultaneously target multiple tumor-promoting signals using GLYTACs via a single, unified mechanism.

## RESULTS AND DISCUSSION

### Synthesis and validation of CI-M6PR ligand

Following the general GLYTAC design outlined in **Figure 1**, we first generated its CI-M6PR-binding glycopolymer ligand, consisting of a *poly*(ethylene glycol) backbone decorated with mannose-6-phosphonate (**M6Pn**) glycans and end-functionalized with biotin (***p*(Man6Pn)-biotin, Figure 2A**) for subsequent conjugation to the glycomimetic heparin arm. The polyvalent display of M6Pn is essential for high-avidity CI-M6PR binding and stabilizing the oligomeric complex.^17,18^ Building on prior work by Banik *et al*.,^16^ who established efficient CI-M6PR-dependent lysosomal uptake of closely related glycopolymers with a degree of polymerization (DP) of ~20, we synthesized *poly*(epichlorohydrin) precursor backbone **P1** of similar length (DP ~ 35) for further elaboration (**Figure 2A**).

**Figure 2.**
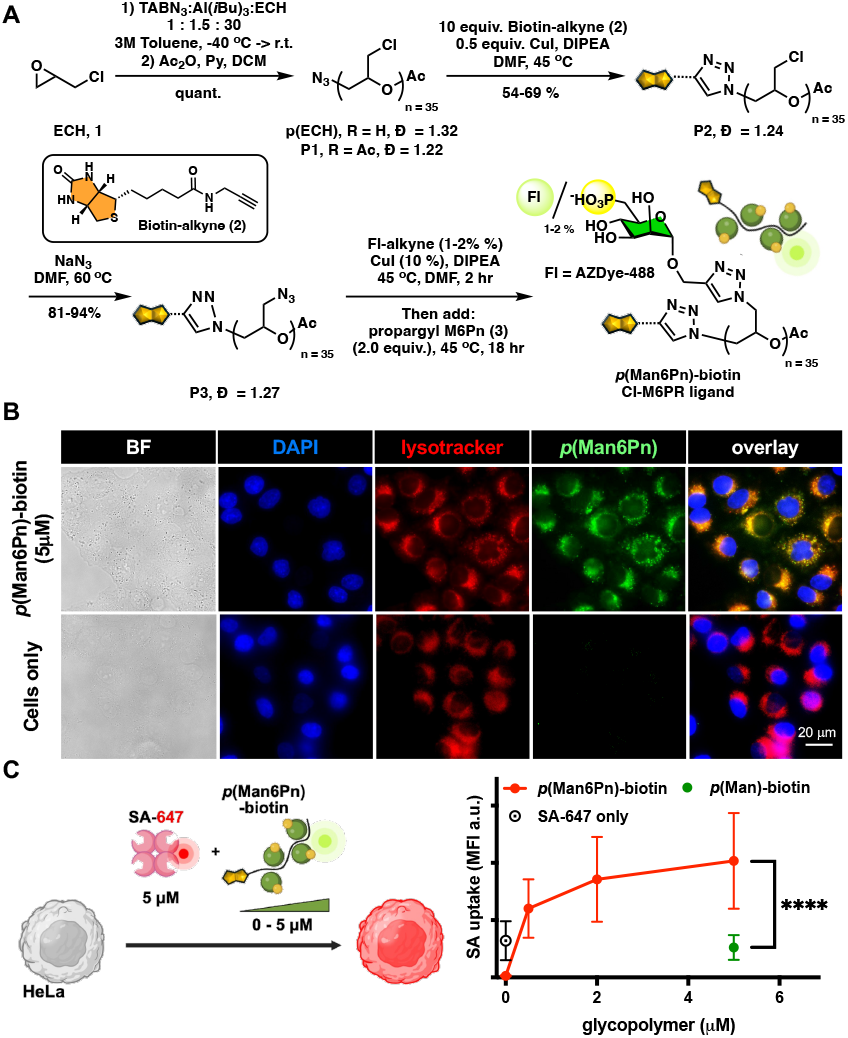
Synthesis and lysosomal targeting of *p*(Man6Pn)-biotin CI-M6PR ligand in HeLa cells. (A) The fluorescently labeled (AZDye-488) and chain-end biotinylated mannose 6-phosphonate (M6Pn) glycopolymer ligand (***p*(Man6Pn)-biotin**) was assembled from a ***p*(ECH)** precursor (DP = 35) through a sequential chain-end functionalization and side-chain glycosylation via the CuAAC reaction. B) Uptake and lysosomal targeting of ***p*(M6Pn)-biotin** (green) by HeLa cells (1 hr, 37°C) via confocal microscopy. LysoTracker (red) was used to mark the lysosomal compartment. (C) Uptake of fluorescent streptavidin-AF647 (**SA-647**, 0.5 µM) in the presence of ***p*(M6Pn)-biotin** (0.5-5.0 µM) or ***p*(Man)-biotin** (5.0 µM) in HeLa cells by flow cytometry. (mean ± STDev, n = 3, **** = p < 0.001).

Controlled monomer-activated ring-opening polymerization of epichlorohydrin (**1**) in the presence of a tetrabutylammonium azide (TBAN_3_) initiator and a triisobutylaluminum (Al(*i*Bu)_3_) activator, according to a method developed by Carlotti,^19^ yielded poly(epichlorohydrin) (***p*(ECH**) with reasonable control over Mw and chain length distribution (Ð = 1.32) The terminal alcohol group was capped by acetylation with acetic anhydride to yield precursor backbone **P1** and to the opposing azide polymer chain end group was appended alkynyl-biotin (**2**) via the CuAAC click reaction^20^ to produce end-functionalized intermediate **P2 (Figure S2**). The chloromethyl sidechains in **P2** were converted to azides in **P3** by treatment with sodium azide (NaN_3_) without a significant increase in chain length dispersity (Ð = 1.27, **Figure 2A**). The assembly of the fluorescent ***p*(Man6Pn)-biotin** CI-M6PR ligand was completed through sequential CuAAC conjugation of AZDye-488-alkyne (~1-2% of azide side chains) and propargyl Man6Pn glycoside (**3, Figure 2A**). The elaboration of the glycopolymer was monitored by IR (ν = 2160 cm^-1^, **Figure S6**), and the resulting ***p*(M6Pn)-biotin** glycopolymer was purified by size exclusion and characterized by ^1^H NMR and UV-Vis spectroscopy (see *Supporting Information*). As a negative control for biological assays, we also generated a glycopolymer analog, ***p*(Man)-biotin**, bearing mannose residues that lack the C6-phosphonate group required for CI-M6PR binding (**Scheme S2**).

Cellular internalization and lysosomal targeting of the glycopolymer CI-M6PR ligand were examined by incubating HeLa cells with AZDye-488–labeled ***p*(M6Pn)-biotin** at increasing concentrations (0.5, 2.0, and 5.0 μM) for 1 h at 37 °C, followed by fluorescence microscopy (**Figure 2B** and **Figures S10** and **S11**). A dose-dependent increase in green fluorescence from the AZDye-488 label, colocalizing with LysoTracker staining (red), demonstrated efficient uptake of ***p*(M6Pn)-biotin** and its accumulation within lysosomes.

Protein internalization via CI-M6PR was further investigated by co-incubating HeLa cells with AlexaFluor647-labeled streptavidin (**SA-647**, 0.5 μM) in the presence of ***p*(M6Pn)-biotin** (0.5-5.0 μM) for 1 h at 37 °C, followed by flow cytometry analysis (**Figure 2C** and **Figure S12**). A significant dose-dependent increase in cellular **SA-647** fluorescence was observed with ***p*(M6Pn)-biotin** compared to untreated controls. In contrast, cells treated with **SA-647** and ***p*(Man)-biotin** (5.0 μM) showed no increase above background, confirming CI M6PR-mediated **SA-647** uptake. At higher concentrations of ***p*(M6Pn)-biotin, SA-647** uptake reached saturation, which we attributed to limited streptavidin availability due to elevated glycopolymer turnover. When dosing **SA-647** at concentrations equimolar to the ***p*(M6Pn)-biotin** ligand (0.5-5.0 μM, **Figure S13**), we observed a linear increase in **SA-647** internalization that scaled with protein availability.

### GLYTAC assembly

Having established ***p*(M6Pn)-biotin** as a suitable vector for lysosomal targeting, we proceeded to elaborate the desired GLYTAC by introducing its growth factor– binding HS-mimetic heparin arm. To connect the two polyanionic GLYTAC domains, we employed a non-covalent streptavidin assembly strategy that we previously developed for the efficient conjugation of heparin-DNA aptamer constructs with defined stoichiometry (**Figure 3A**).^21^

**Figure 3.**
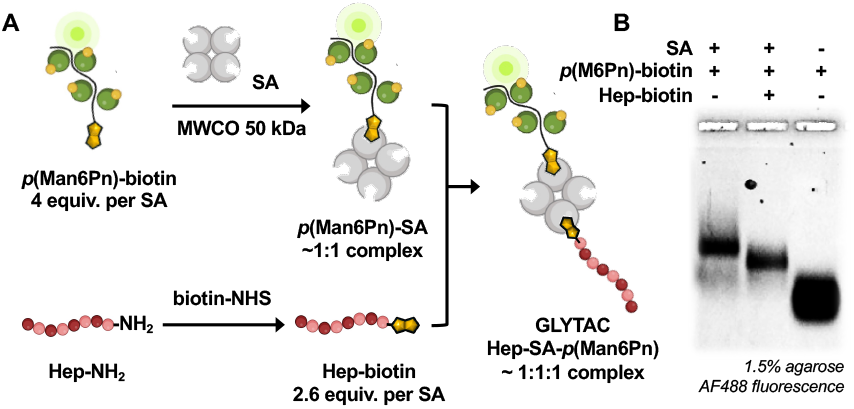
Assembly of Hep-SA-*p*(Man6Pn) GLYTAC conjugate. (A) The **GLYTAC** was assembled by sequential addition of ***p*(M6Pn)-biot**in to a **SA**, followed by addition of **Hep-biotin**, with 50 kDa spin filtration at each step to remove unbound components and yield an equimolar **Hep-SA-*p*(M6Pn)** conjugate. (B) Native agarose gel electrophoresis (1.5%) visualized by AZDye-488 fluorescence showed reduced mobility of the ***p*(M6Pn)-SA** complex relative to ***p*(M6Pn)-biotin**, and a further shift upon **Hep-biotin** incorporation, consistent with the addition of the negatively charged heparin arm.

Accordingly, tetrameric **SA** (16 μM) was incubated with increasing equivalents of ***p*(M6Pn)-biotin** (0.5 - 5.0 equiv.) for 24 h and the efficiency of complex formation was assessed by native agarose gel electrophoresis (1.5%, **Figure S8**). The binding of **SA** to ***p*(M6Pn)-biotin** produced a new, increasingly fluorescent band corresponding to ***p*(M6Pn)-SA**, which showed reduced electrophoretic mobility on the gel compared to the parent glycopolymer. Treating **SA** with 4 equivalents of the biotinylated glycopolymer maximized the formation of a 1:1 ***p*(M6Pn)-SA** complex while preventing higher stoichiometries, as confirmed by UV–Vis absorption measurements at λ = 280 nm for **SA** and 495 nm for ***p*(M6Pn)** after purification by spin filtration (MWCO 50 kDa) to remove the unbound constituents **Figures 3B** and **S9**).

Further treatment of the ***p*(M6Pn)-SA** complex (11 μM) with **Hep-biotin**^21^ (29 μM, 2.6 equiv per **SA**) in DPBS buffer (pH 7.4, 24 h) yielded the desired **Hep-SA-*p*(M6Pn)** conjugate (**Figure 3A**). Native agarose gel electrophoresis (1.5%) of the purified **GLYTAC**, following removal of residual components by spin filtration (MWCO 50 kDa), showed a slight band shift toward higher electrophoretic mobility, consistent with incorporation of the highly negatively charged heparin polysaccharide chain (**Figure 3B**). To determine complex stoichiometry, we digested some of the **GLYTAC** conjugate with heparinases I–III and analyzed HS disaccharides by glycan reductive isotope labeling (GRIL) LTQ-MS (**Table S1**). Normalization to protein content by BCA assay established an approximate 1:1 **Hep-biotin** to **SA** ratio, confirming an overall near 1:1:1 **Hep-SA-*p*(M6Pn) GLYTAC** stoichiometry. Increasing **Hep-biotin** equivalents did not produce higher-stoichiometry GLYTAC assemblies, presumably due to steric hindrance and charge repulsion between the macromolecular **Hep** and ***p*(M6Pn)** arms. Notably, reversing the assembly order by adding ***p*(M6Pn)-biotin** to a preformed **Hep-SA** (1:1) complex was unsuccessful, likely due to limited accessibility of the biotin handle positioned without a spacer and directly adjacent to the negatively charged ***p*(M6Pn)** glycodomain.

### HS-mimetic GLYTAC mediates lysosomal clearance of extracellular FGF2

Using FGF2 as a prototypical extracellular HS-binding growth factor associated with enhanced tomor cell proliferation and resistance to anti-VEGF therapies, we assessed the CI-M6PR-dependent uptake and lysosomal degradation of the protein in the presence of the HS-mimetic **GLYTAC** (**Figure 4A**). HeLa cells were incubated with FGF2 (200 nM) and **Hep-SA-*p*(M6Pn)** (2 μM, DMEM with 10% FBS, 37 °C), and protein internalization was monitored by immunostaining and confocal fluorescence microscopy over 60 min (**Figure 4B** and **Figure S14**). Cells treated with FGF2 alone or with **Hep-SA** (2 μM) lacking the CI-M6PR ligand arm served as controls. Lysosomes were delineated by LAMP2 immunostaining. In cells treated with FGF2 alone, the protein localized as fluorescent puncta at or near the cell surface within 5 min, then internalized into intracellular regions devoid of LAMP2 signal, consistent with endocytosis mediated by endogenous HSPGs ^22^(**Figure S12**). Inclusion of **Hep-SA** produced robust and persistent FGF2 staining at the cell surface and intracellularly, outside lysosomes, indicating protein stabilization and inhibition (**Figure 4**). By contrast, **Hep-SA-*p*(M6Pn)** promoted rapid and uniform internalization of FGF2 within 5 min, with complete overlap of the FGF2 and LAMP2 signals at 30 min and near disappearance of FGF2 by 60 min, consistent with lysosomal targeting and degradation (**Figure 4**). Addition of the broad-spectrum protease inhibitor Leupeptin (0.1 mg/mL) blocked lysosomal degradation, leading to accumulation of FGF2 within the LAMP2-positive regions, further confirming **GLYTAC**-mediated lysosomal clearance of the growth factor (**Figure 4**).

**Figure 4.**
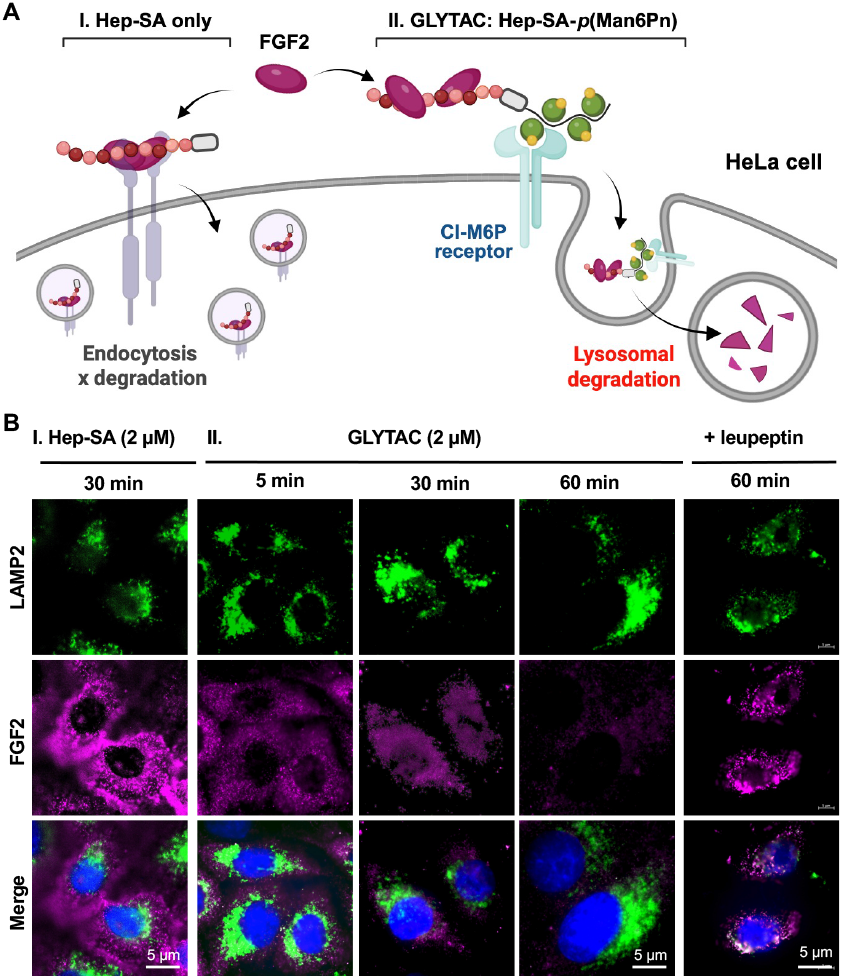
HS-mimetic GLYTAC mediates lysosomal clearance of extracellular FGF2 in HeLa cells. (A) Differential uptake and degradation of FGF2 with **Hep-SA** or **Hep-SA-*p*(M6Pn)**. (B) Confocal fluorescence micrographs of HeLa cells incubated with FGF2 (200 nM) ± **Hep-SA** or **Hep-SA-*p*(M6Pn)** (2 μM) for 5-60 min. FGF2 localization was assessed by immunostaining, with LAMP2 antibody marking lysosomes. **Hep-SA** stabilized FGF2 outside lysosomes, whereas **Hep-SA-*p*(M6Pn)** induced rapid internalization, colocalization with LAMP2, and subsequent degradation. Leupeptin (0.1 mg/mL) blocked degradation, leading to FGF2 accumulation in LAMP2-positive compartments.

### HS-mimetic GLYTACs enable targeted degradation of FGF receptors

Cell-surface HS facilitates assembly of FGF2 into a functional signaling complex with its receptors (FGFRs). Although most FGFRs (i.e., types 1–3) bind HS only weakly, they associate strongly with the FGF2–HS complex, enabling internalization and downstream signaling.^13^ This raises the possibility that GLYTACs could target not only FGF2 but the entire signaling complex for degradation, resulting in sustained suppression of FGFR-mediated signaling.

To test this hypothesis, serum-starved HeLa cells were treated with FGF2 (20 ng/mL) in the presence of **Hep-SA-*p*(M6Pn)** (0.2 nM to 2.0 μM) for 2 hrs at 37 °C. Cells exposed to FGF2 alone, or in the presence of **Hep-SA**, across the same concentration range, served as controls. Following treatment, cells were lysed, and FGFR1 levels were assessed by Western blot analysis (**Figure 5**). **HepSA** produced a dose-dependent increase in FGFR1 levels after FGF2 stimulation (**Figure 5B**), consistent with the observed stabilization of FGF2 by this conjugate following internalization (**Figure 3A**). At lower concentrations (0.2 – 20 nM), **Hep-SA-p(M6Pn)** stabilized FGFR1 similarly to **Hep-SA**; however, at concentrations above 200 nM the **GLYTAC** induced FGFR1 degradation to levels similar to FGF2-unstimulated cells (**Figure 5B**). This shift from stabilization to degradation indicates CI-M6PR engagement, consistent with the EC50 of ~ 430 nM measured for **Hep-SA-*p*(M6Pn)**-mediated **SA-647** uptake in HeLa cells (**Figure 2C**).

**Figure 5.**
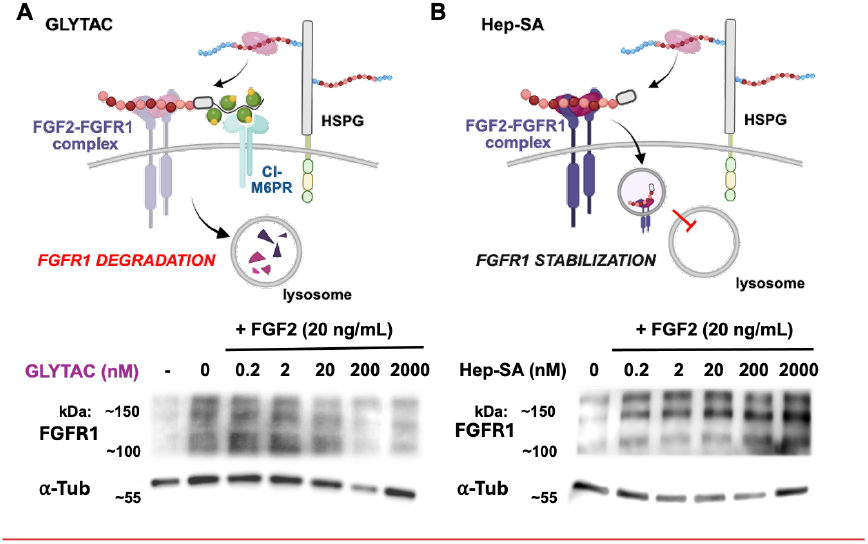
HS-mimetic GLYTAC promotes FGFR1 degradation in HeLa cells. Serum-starved HeLa cells were stimulated with FGF2 (20 ng/mL) and (A) **Hep-SA-*p*(M6Pn)** or (B) **Hep-SA** (0.2-2000 nM) for 2 hrs at 37 °C. FGFR1 levels were analyzed by Western blot with α-Tubulin as a loading control. **Hep-SA** stabilized FGFR1, whereas **Hep-SA-*p*(M6Pn)** induced degradation above 200 nM, consistent with CI-M6PR engagement.

Collectively, our results indicate that HS-mimetic GLYTACs can enforce internalization and lysosomal degradation of both extracellular FGF2 and cell surface FGFR1, providing a potential mechanism for sustained therapeutic suppression of growth factor signaling in the tumor environment.

## CONCLUSIONS

The tumor microenvironment is complex, with cancer cells exploiting diverse signals to sustain survival. LYTACs, chemical degraders of extracellular proteins that harness lysosomal recycling pathways, have emerged as a promising new class of cancer therapeutics.^16,23^ However, current LYTAC architectures, typically based on specific antibodies, are limited in the number of proteins they can simultaneously target. Expanding LYTACs to eliminate multiple pro-tumorigenic cues through a shared molecular feature could enable more effective combination therapies. Many growth factors driving tumor progression – including FGF, VEGF, and TGF-β – require binding to cell-surface HS glycans for activity.^13,14,15^ This shared dependence provides an opportunity to exploit HS interactions for targeted lysosomal clearance. In this study, we report a prototype HS-glycomimetic degrader (GLYTAC) and demonstrate its activity in HeLa cells. By linking heparin, a prototypical HS mimetic, to a poly(M6Pn) ligand for CI-M6PR-mediated uptake, we show rapid internalization and degradation of FGF2. Owing to the enhanced affinity of FGFRs for HS-bound FGF2, GLYTAC treatment also promoted FGFR1 degradation, further reducing the availability of pro-tumorigenic signals in the cancer cell environment. Ongoing work focuses on optimizing GLYTAC components to enhance efficiency and specificity, with the ultimate goal of achieving durable suppression of growth factor signaling and reduced cancer cell survival, particularly in drug-resistant tumors.

## ASSOCIATED CONTENT

### Supporting Information

Specifications for all reagents and instrumentation, synthetic procedures and structural characterization for synthetic precursors and bioconjugates, detailed protocols for biological experiments, and expanded experimental data for **Figures 2-5** are included in the Supporting Information, which is available free of charge on the ACS Publications website.

## AUTHOR INFORMATION

### Author Contributions

The manuscript was written through the contributions of all authors. K.G. and J.L.F. designed the research; J.L.F. and T.T. performed experiments; J.L.F., T.T., and K.G. analyzed data; J.L.F. and K.G. wrote the manuscript and prepared the supporting information. All authors have given approval to the final version of the manuscript.

## ACKNOWLEDGMENT

This work was supported by grant 5R01GM145913-02 (NIGMS). We thank the UCSD Microscopy Core Facility for assistance with fluorescence microscopy (via NINDS P30 Grant: P30NS047101 and S10 Grant: S10OD030505) and the UCSD Glycobiology Research and Training Center. We thank Dr. Shubham Parashar for assistance with FGFR Western blotting assays, Drs. Esko and Gordts for valuable discussions, and Dr. Ryan Porell for contributions to early concept development.

