## Supplementary information for "Glycomimetic Lysosome-targeting Chimeras (GLYTACs) for Degradation of Growth Factors and Growth Factor Receptors in Cancer Cells"

### TABLE OF CONTENTS


**ABBREVIATIONS** ..... **S4**

**MATERIALS AND REAGENTS** ..... **S5**

**INSTRUMENTATION** ..... **S6**

**GENERAL SYNTHETIC CHEMISTRY PROCEDURES** ..... **S7**

SYNTHESIS OF PROPARGYL MANNOSE 6-PHOSPHONATE (**MAN6PN**) GLYCOSIDE **3**. ..... S8

*Scheme S1. Synthesis of propargyl mannose-6-phosphonate glycoside 3.* ..... *S8*

*Synthesis of benzyl α-D-mannopyranoside (S-1).* ..... *S8*

*Synthesis of benzyl 2, 3, 4, 6-tetra-O-trimethylsilylmanoside (S-2).* ..... *S9*

*Synthesis of benzyl 2,3,4-tri-O-trimethylsilylmanoside (S-3).* ..... *S9*

*Synthesis of diethyl ((E)-2-((2R,3R,4S,5S,6S)-6-(benzyloxy)-3,4,5-tris((trimethylsilyl)oxy)-tetrahydro-2H-pyran-2-yl)vinyl)phosphonate (S-4).* ..... *S10*

*Synthesis of (2S,3S,4S,5R,6R)-2-(benzyloxy)-6-((E)-2-(diethoxyphosphoryl)vinyl) tetrahydro -2H-pyran-3,4,5-triyl triacetate (S-5).* ..... *S11*

*Synthesis of (2R,3R,4S,5S)-2-((E)-2-(diethoxyphosphoryl)vinyl)-6-hydroxytetrahydro-2H-pyran-3,4,5-triyl triacetate (S-6).* ..... *S12*

*Synthesis of (2R,3R,4S,5S)-2-((E)-2-(diethoxyphosphoryl)vinyl)-6-(2,2,2-trichloro-1- imino-ethoxy)tetrahydro-2H-pyran-3,4,5-triyl triacetate (S-7).* ..... *S13*

*Synthesis of (2-((2R,3R,4S,5S,6S)-3,4-diacetoxy-6-(prop-2-yn-1-yloxy)-5-((trimethylsilyl)-oxy)tetrahydro-2H-pyran-2-yl)ethyl)phosphonic acid (S-8).* ..... *S13*

*Synthesis of propargyl mannose 6-phosphonate glycoside (3).* ..... *S14*

SYNTHESIS OF GLYCOPOLYMERS ***P*(MAN6PN)-BIOTIN** AND ***P*(MAN)-BIOTIN**. ..... S15

*Scheme S2. Synthesis of glycopolymers p(Man6Pn)-biotin and p(Man)-biotin.* ..... *S15*

*Synthesis of chain-end O-acetylated poly(epichlorohydrin) backbone P1.* ..... *S15*

*Figure S1. GPC trace of chain-end O-acetylated poly(epichlorohydrin) backbone P1* ..... *S16*

*Synthesis of chain-end-biotinylated poly(epichlorohydrin) polymer P2.* ..... *S16*

*Figure S2. GPC trace of chain-end biotinylated poly(epichlorohydrin) polymer P2.* ..... *S17*

*Figure S3. IR spectrum of chain-end biotinylated poly(epichlorohydrin) polymer P2.* ..... *S17*

*Synthesis of chain-end biotinylated poly(glycidyl azide) polymer P3.* ..... *S17*

*Figure S4. GPC trace of chain-end biotinylated poly(glycidyl azide) polymer P3.* ..... *S18*

*Figure S5. IR spectrum of chain-end biotinylated poly(glycidyl azide) polymer P3.* ..... *S18*

*Synthesis of CI-M6PR glycopolymer ligand p(M6Pn)-biotin.* ..... *S18*

*Figure S6. IR spectrum of CI-M6PR glycopolymer ligand p(M6Pn)-biotin.* ..... *S20*

*Synthesis of CI-M6PR non-binding glycopolymer p(Man)-biotin.* ..... *S20*

*Figure S7. IR spectrum of CI-M6PR non-binding glycopolymer p(Man)-biotin.* ..... *S21*

GLYTAC ASSEMBLY AND CHARACTERIZATION ..... S21
*Reducing end-biotinylation of heparin (**Hep-biotin**)* ..... S21
*Assembly of **p(M6Pn)-SA** 1:1 complex.* ..... S22
***Figure S9.** Analysis of purified **p(M6Pn)-SA** conjugate.* ..... S24
*Assembly of **Hep-SA-p(Man6Pn)** GLYTAC conjugate* ..... S24
*Determination of hep/SA complex stoichiometry via (GRIL) LTQ–MS disaccharide analysis* ..... S25
**BIOLOGICAL PROCEDURES ..... S26**
CELL CULTURE ..... S26
INTERNALIZATION AND LYSOSOMAL TARGETING OF *P*(M6PN) IN HELA CELLS BY MICROSCOPY. ..... S26
***Figure S10.** Internalization of **p(M6Pn)*** ..... S27
***Figure S11**. Dose-dependent internalization of **p(M6Pn)*** ..... S28
*P*(M6PN)-MEDIATED SA647 UPTAKE BY FLOW CYTOMETRY. ..... S28
***Figure S12**. Uptake of **SA-647** in HeLa cells mediated by equimolar **p(M6Pn)-biotin**.* ..... S29
***Figure S13.** Uptake of **SA-647** in HeLa cells mediated by equimolar **p(M6Pn)-biotin**.* ..... S29
FGF2 DEGRADATION IMMUNOHISTOCHEMISTRY (IHC). ..... S30
***Figure S14**. Internalization of FGF2 in HeLa cells (expanded data for **Figure 4B**).* ..... S31
FGFR DEGRADATION BY WESTERN BLOTTING ..... S31
***Figure S15**. Western blot analysis of FGFR1 levels in HeLa cells.* ..... S31
**REFERENCES ..... S34**
**APPENDIX: $^{1}$H AND $^{13}$C NMR SPECTRA OF SYNTHETIC COMPOUNDS ..... S35**

#### ABBREVIATIONS

**AF488** = AlexaFlour 488

**AF647** = AlexaFlour 647

**BCA** = bicinchoninic acid

**BSA** = bovine serum albumin

**DCM** = dichloromethane

**DMSO** = dimethyl sulfoxide

**DMEM** = Dulbecco's modified Eagle medium

**DMF** = N,N-dimethylformamide

**DPBS** = Dulbecco's phosphate buffer saline

**ESI-MS** = electrospray ionization mass spectrometry

**EtOAc** = ethyl acetate

**FBS** = fetal bovine serum

**FGF** = fibroblast growth factor

**GLYTAC** = glycomimetic lysosome-targeting chimera

**GP** = glycopolymer

**GPC** = gel permeation chromatography

**HCl** = hydrochloric acid

**Hep** = heparin

**Hex** = n-hexanes

**HRP** = horseradish peroxidase

**HS** = heparan sulfate

**HSPG** = heparan sulfate proteoglycan

**IR** = infrared spectroscopy

**IHC** = immunohistochemistry

**LYTAC** = lysosome-targeting chimera

**M6P** = Mannose 6-phosphate

**M6Pn** = Mannose 6-phosphonate

**M6PR** = Mannose 6-phosphate receptor

**MALDI-TOF** = matrix-assisted laser desorption/ionization – time of flight

**MQ** = milli-Q ultrapure water

**MWCO** = molecular weight cut-off

**NHS** = N-hydroxysuccinimide

**NMR** = nuclear magnetic resonance

**PBS** = phosphate buffered saline

**PD-10** = prepacked sephadex® G-25 medium disposable column

**PG** = proteoglycan

**RBF** = round-bottom flask

**TFA** = trifluoroacetic acid

**THF** = tetrahydrofuran

**TLC** = thin-layer chromatography

**TMSCl** = trimethylsilyl chloride

#### MATERIALS AND REAGENTS

| Chemical Reagents | Source | Catalog No. |
|---|---|---|
| 1,8-diazabicyclo [5.4.0]undec-7-ene (DBU) | Sigma Aldrich | 139009-25G |
| 2-aminoacridone | Sigma Aldrich | 6627 |
| 3-azido-7-hydroxycoumarin | Carbosynth | FA31762 |
| Biotin-dPEG$_{11}$-Azide | Quanta Biodesign | 10784 |
| Carbazole | Ultra Scientific | HAH-022 |
| NHS-BCN | Sigma Aldrich | 744867-10MG |
| TAMRA-NHS | Invitrogen | C300 |
| Triton X-100 | Alfa Aesar | A16046 |
| Tween-20 | VWR | M147-4L |
| TMB substrate solution | VWR | 97063-666 |
| Phosphate Buffered Saline with Ca/Mg | Corning | 21-030-CM |
| **Biological Reagents** | **Source** | **Catalog No.** |
| Bovine Serum Albumin (BSA) | Spectrum | A3611 |
| Fibroblast growth factor 2 | Peprotech | 100-18B |
| Anti-FGF2 | Millipore | 05-118 |
| Anti-FGFR1 | Cell Signaling | 9740S |
| Anti-LAMP2-AF555 | Invitrogen | MA1-205-A555 |
| Anti-mouse IgG HRP | Cell Signaling | 7076S |
| Anti-rabbit IgG HRP | Cell Signaling | 7074S |
| Heparin | Iduron | HEP001 |
| Streptavidin | Leinco | S203 |
| Streptavidin AF647 | Invitrogen | S21374 |

| Consumables | Source | Catalog No. |
|---|---|---|
| 3 kDa Centrifugal Spin Filters | Amicon | UFC500396 |
| 50 kDa Centrifugal Spin Filters | Amicon | UFC505096 |
| 1.5 mL Microcentrifuge Tube | Thermo Fisher | 5408129 |
| HEPES | Gibco | 15630106 |
| PCR tube | Thermo Fisher | AB0337 |
| PD-10 Desalting Column | GE Life Sciences | 17085101 |
| Trypsin-EDTA (0.25%) | Gibco | 25200114 |

#### INSTRUMENTATION

Flash chromatography was performed on a Biotage Isolera One automated flash chromatography system. All NMR spectra were recorded on Joel AV 500 MHz NMR spectrometer. High-resolution mass spectra (HRMS), ESI mode, were obtained on an Agilent 6230 ESI-TOFMS in positive ion mode at the Molecular Mass Spectroscopy Facility at the Chemistry and Biochemistry Department, University of California San Diego. UV-Vis spectra for polymer fluorophore content quantification were collected using a quartz cuvette using a Thermo Scientific Nanodrop2000c spectrophotometer. IR spectroscopy was performed on a Nicolet 6700 FT-IR spectrophotometer (Thermo Scientific). Size exclusion chromatography (SEC) was performed on a Hitachi Chromaster system equipped with an RI detector and two 5 µm, mixed bed, 7.8 mm I.D. x 30 cm TSKgel columns in series (Tosoh Bioscience). Absorbance values were measured on ThermoFisher NanoDrop One. Fluorescence microscopy was performed on Zeiss Elyra 7 Lattice microscopy. Images were analyzed using Zeiss ZEN software and ImageJ. Flow cytometry analysis was performed on BD Accuri C6 flow cytometer and analyzed with FlowJo software. Gels and membranes were imaged using Amersham A680 RGB imager and analyzed with ImageJ. Data was exported and analyzed in GraphPad Prism 10 software.

#### GENERAL SYNTHETIC CHEMISTRY PROCEDURES

Poly(epichlorohydrin) precursor backbone ***p*(ECH)** (Mn = 3056, PD = 35, Đ = 1.32) was prepared by controlled polymerization of epichlorohydrin according to a method by Carlotti,[1] as reported previously by Honigfort *et al.*[2] Unless otherwise noted, all reagents were used as received and reactions were performed in oven- or flame-dried glassware under an atmosphere of nitrogen equipped with rubber septa, and magnetic stirring. All solvents were anhydrous and transferred via stainless steel syringe. Reactions were monitored by electrospray ionization liquid chromatography-mass spectrometry (ESI LC-MS) and thin layer chromatography (TLC) in which glass plates coated with silica gel were used and visualized with shortwave 254 nm UV light and/or developed upon heating with p-anisaldehyde (PAA) or potassium permanganate ($KMNO_4$). Proton chemical shifts were recorded in parts per million (ppm) on the δ scale, downfield from tetramethylsilane and referenced from an internal standard of the residual protium in the NMR solvents. Data for 13C NMR were reported in ppm downfield from tetramethylsilane and referenced based on the chemical shift from the carbon resonances of the solvent. NMR data were reported as follows: chemical shift, multiplicity (s = singlet, d = doublet, t = triplet, q = quartet, m = multiplet, br = broad), coupling constant in Hz, integration. Organic soluble polymers were analyzed using an isocratic method with a flow rate of 0.7 mL/min in DMF (0.2% LiBr, 70 °C). Intermediates **S-3** through **S-7** were synthesized in accordance to previously published methods and NMR data matched those published.[3]

#### Synthesis of propargyl mannose 6-Phosphonate (Man6Pn) glycoside 3.

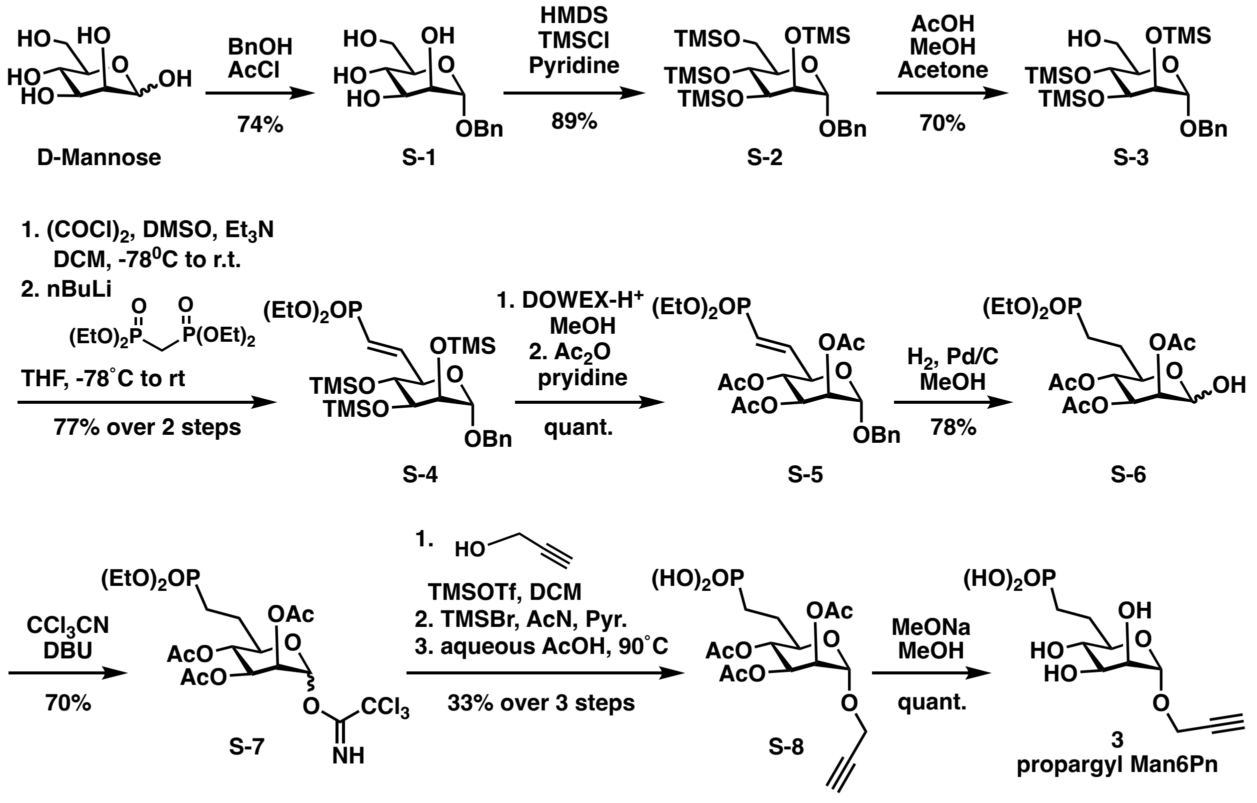


**Scheme S1.** Synthesis of propargyl mannose-6-phosphonate glycoside **3**.

##### Synthesis of benzyl α-D-mannopyranoside (S-1).

D-(+)-Mannose (6.53 g) was added to a dried round-bottom flask equipped with a magnetic stir bar. The flask was fitted with a septum and flushed with $N_2$. Benzyl alcohol (30.0 mL) was added, and the suspension was heated to 50 °C with stirring. Acetyl chloride was added dropwise over 15 min. After 1 h, the reaction was cooled to room temperature and stirred for an additional 19 h. Reaction progress was monitored by TLC (5% MeOH/DCM). The mixture was carefully poured into deionized water (100 mL) and extracted with ethyl acetate (3 × 50 mL). The combined organic layers were washed with deionized water (2 × 50 mL), and the aqueous phases were collected and concentrated with toluene. The resulting oil was purified by flash column chromatography (3 - 10% MeOH/DCM) to afford benzyl mannoside **S-1** as a white solid (7.26 g, 74%). $^1$H NMR (500

MHz, $CD_3OD$) δ: 7.33 – 7.23 (m, 6H), 4.80 (d, $J$ = 1.7 Hz, 1H), 4.71 (dd, $J$ = 11.8, 1.9 Hz, 1H), 4.47 (dd, $J$ = 11.8, 1.9 Hz, 1H), 3.83 (t, $J$ = 2.0 Hz, 1H), 3.81 – 3.79 (m, 2H), 3.71 (q, $J$ = 1.9 Hz, 1H), 3.70 – 3.67 (m, 2H), 3.61 (dd, $J$ = 9.8, 1.9 Hz, 1H), 3.59 – 3.55 (m, 1H). $^{13}$C NMR (126 MHz, $CD_3OD$) δ: 38.96, 129.38, 128.76, 100.61, 74.82, 72.59, 72.15, 69.84, 62.87. HRMS (ESI-Pos) *m/z* calculated for $C_{13}H_{18}O_6$ (270.11): Found 293.0996 $[M+Na]^+$.

**Synthesis of benzyl 2, 3, 4, 6-tetra-*O*-trimethylsilylmanoside (S-2).**

Benzyl α-D-mannopyranoside (**S-1**, 2.30 g) and a magnetic stir bar were added to a dried 100 mL round-bottom flask. The flask was flushed with $N_2$, and pyridine (35.0 mL) was added. The solution was stirred at 0 °C. A solution of TMSCl (6.50 mL) in HMDS (3.60 mL) was added dropwise. Upon complete addition, the reaction mixture was gradually warmed to room temperature and stirred under inert atmosphere overnight. After 20 h, no starting material was detected by TLC (5% MeOH/DCM). The solvent was removed under reduced pressure. The resulting white solid was suspended in pentane and extracted with deionized water (2 × 30 mL). The aqueous layers were back-washed with pentane (2 × 20 mL), and the organic layers were combined, dried over anhydrous magnesium sulfate, and concentrated *in vacuo* to afford **S-2** as a clear oil (4.23 g, 89%). $^1$H NMR (500 MHz, $CDCl_3$) δ: 7.33 (m, 5H, aromatic), 4.73 (d, $J$ = 12.1 Hz, 1H), 4.67 (d, $J$ = 1.7 Hz, 1H), 4.47 (d, $J$ = 12.1 Hz, 1H), 3.92 – 3.69 (m, 6H), 0.16 (s, 9H), 0.15 (s, 9H), 0.13 (s, 9H), 0.10 (s, 9H). $^{13}$C NMR (126 MHz, $CDCl_3$) δ 138.18, 128.50, 127.85, 127.70, 77.50, 77.25, 76.99, 75.16, 73.84, 73.04, 68.71, 68.50, 62.69, 22.55, 14.26, 0.89, 0.75, 0.59, 0.11. HRMS (ESI-Pos) *m/z* calculated for $C_{25}H_{50}O_6Si_4$ (558.27): Found 509.22 $[M\text{-}6'TMS+Na]^+$.

**Synthesis of benzyl 2,3,4-tri-O-trimethylsilylmanoside (S-3).**

Benzyl 2,3,4,6-tetra-O-trimethylsilylmannoside (**S-2**, 2.34 g, 4.19 mmol) was weighed into an oven-dried 100 mL round-bottom flask equipped with a magnetic stir bar. The flask was fitted with a septum and flushed with Ar. Methanol (13.5 mL) and acetone (9.20 mL) were added, and the solution was stirred at 0 °C for 15 min. Acetic acid (0.500 mL, 8.38 mmol, 2.00 equiv) was added dropwise. The reaction mixture was stirred at 0 °C for 4.5 h and monitored by TLC (10% EtOAc/hexanes). The reaction was quenched by addition of solid sodium bicarbonate (0.900 g). The suspension was stirred at ambient temperature for 10 min and then concentrated *in vacuo*. The crude residue was purified by flash silica gel chromatography (0–15% EtOAc/hexanes) to yield **S-3** as a pale, tan-colored translucent oil (1.43 g, 70%). $^1$H NMR (500 MHz, $CDCl_3$) δ: 7.39 – 7.26 (m, 5H, aromatic), 4.71 (d, *J* = 12.2 Hz, 1H), 4.69 (s, 1H), 4.49 (d, *J* = 12.1 Hz, 1H), 3.89 (t, *J* = 8.8 Hz, 1H), 3.85 – 3.78 (m, 2H), 3.75 (dd, *J* = 11.5, 2.9 Hz, 1H), 3.69 (dd, *J* = 11.5, 5.2 Hz, 1H), 3.64 – 3.55 (m, 1H), 0.17 (s, 9H), 0.15 (s, 9H), 0.11 (s, 9H). HRMS (ESI-Pos) *m/z* calculated for $C_{22}H_{42}O_6Si_3$ (486.23): Found 509.2182 $[M+Na]^+$.

**Synthesis of diethyl ((*E*)-2-((2*R*,3*R*,4*S*,5*S*,6*S*)-6-(benzyloxy)-3,4,5-tris((trimethylsilyl)oxy)-tetrahydro-2*H*-pyran-2-yl)vinyl)phosphonate (S-4).**

An oven-dried round-bottom flask was fitted with a magnetic stir bar and septum, then flushed with $N_2$. Oxalyl chloride (108 µL, 1.25 mmol, 1.10 equiv) and dichloromethane (2.00 mL) were added, and the mixture was stirred at –78 °C. DMSO (180 µL, 2.51 mmol, 2.20 equiv) was added dropwise. After 10 min, a solution of benzyl 2,3,4-tri-O-trimethylsilylmannoside (**S-3**, 0.555 g, 1.14 mmol) in DCM (1.50 mL) was added dropwise over several minutes. The reaction was stirred at –78 °C under $N_2$ for 45 min, then triethylamine (800 µL, 5.70 mmol, 5.00 equiv) was added slowly. The flask was warmed to ambient temperature and stirred for an additional 1 h. Reaction progress was monitored by TLC (10% EtOAc/hexanes). The mixture was diluted with DCM and washed with deionized $H_2O$ (3 × 10 mL). The organic phase was dried over anhydrous magnesium sulfate, filtered, and concentrated. The resulting pale yellow oil was immediately carried forward. A dried round-bottom flask was charged with tetraethyl methylene diphosphate (425 µL, 1.71 mmol, 1.50 equiv) and THF (2.00 mL). The solution was cooled to –78 °C, and n-BuLi (2.5 M in hexanes, 0.600 mL, 1.43 mmol, 1.25 equiv) was added dropwise. After stirring under $N_2$ for 45 min, a THF solution of the intermediate (1.50 mL) was added slowly. The reaction was stirred overnight at ambient temperature. Reaction progress was monitored by TLC (10% EtOAc/hexanes). The pale yellow mixture was diluted with EtOAc and washed with deionized $H_2O$ (2 × 10 mL). The aqueous layer was extracted with EtOAc, and the combined organic phases were dried over anhydrous magnesium sulfate, filtered, and concentrated to yield vinyl phosphonate **S-4** as a tan oil (0.540 g, 77%). $^1$H NMR (500 MHz, $CDCl_3$) δ: 7.37 – 7.21 (m, 5H), 6.89 – 6.77 (m, 1H), 6.09 – 5.95 (m, 1H), 4.69 (t, *J* = 2.3 Hz, 1H), 4.67 – 4.60 (m, 1H), 4.47 – 4.41 (m, 1H), 4.08 (ddt, *J* = 11.0, 9.4, 4.7 Hz, 5H), 3.84 – 3.75 (m, 2H), 3.70 – 3.63 (m, 1H), 1.34 – 1.19 (m, 6H), 0.19 – 0.03 (m, 27H). $^{13}$C NMR (126 MHz, $CDCl_3$) δ: 171.17, 149.04, 137.61, 129.16 – 126.97, 117.81, 116.31, 100.30, 74.26 – 72.66, 72.89, 71.55, 69.17, 61.74, 60.43, 21.09, 16.48, 14.26, 1.07. HRMS (ESI-Pos) *m/z* calculated for $C_{27}H_{51}O_8PSi_3$ (618.26): Found 619.2702 $[M+H]^+$.

**Synthesis of (2*S*,3*S*,4*S*,5*R*,6*R*)-2-(benzyloxy)-6-((*E*)-2-(diethoxyphosphoryl)vinyl) tetrahydro -2*H*-pyran-3,4,5-triyl triacetate (S-5).**

A vial was charged with a magnetic stir bar, vinyl phosphonate **S-4** (0.473 g, 0.765 mmol), and DOWEX 50WX8 ($H^+$ form, 200–400 mesh, 0.360 g). The reaction mixture was suspended in methanol (8.0 mL) and stirred at ambient temperature. Reaction progress was monitored by TLC (50% EtOAc/hexanes) and confirmed complete by mass spectrometry after 2 h. The suspension was filtered into an oven-dried round-bottom flask, and methanol was removed *in vacuo*. The crude material was immediately carried forward by the addition of acetic anhydride (2.0 mL) and pyridine (4.0 mL). The reaction was stirred overnight at ambient temperature under $N_2$. Completion was confirmed by TLC (10% MeOH/DCM) after 16 h. Solvent was removed *in vacuo*, and the crude tan oil was diluted with DCM (10 mL), washed with 1 M HCl (20 mL) and deionized $H_2O$ (20 mL). The aqueous phases were back-extracted with DCM (10 mL), and the combined organic layers were dried over anhydrous magnesium sulfate, filtered, and concentrated to afford **S-5** as a tan oil (0.401 g, 99%). $^1H$ NMR (500 MHz, $CDCl_3$) δ: 7.39 – 7.27 (m, 5H), 6.59 (tt, *J* = 17.1, 5.1 Hz, 1H), 6.04 – 5.93 (m, 1H), 5.39 (dd, *J* = 10.0, 3.4 Hz, 1H), 5.31 – 5.26 (m, 1H), 5.12 (t, *J* = 10.1 Hz, 1H), 4.89 (d, *J* = 1.8 Hz, 1H), 4.68 (d, *J* = 11.9 Hz, 1H), 4.55 (d, *J* = 12.0 Hz, 1H), 4.33 (ddq, *J* = 9.9, 4.9, 1.9 Hz, 1H), 4.16 – 4.02 (m, 4H), 2.13 (s, 3H), 2.02 (s, 3H), 1.97 (s, 3H), 1.33 (td, *J* = 7.1, 1.0 Hz, 6H). HRMS (ESI-Pos) *m/z* calculated for $C_{24}H_{33}O_{11}P$ (528.18): Found 529.1831 $[M+H]^+$.

**Synthesis of (2*R*,3*R*,4*S*,5*S*)-2-((*E*)-2-(diethoxyphosphoryl)vinyl)-6-hydroxytetrahydro-2*H*-pyran-3,4,5-triyl triacetate (S-6).**

An oven-dried round-bottom flask equipped with a magnetic stir bar was charged with **S-5** (0.114 g, 0.215 mmol) dissolved in methanol (4.0 mL). The reaction vessel was flushed with $N_2$, and 10% Pd/C (53.0 mg, 0.0430 mmol, 0.20 equiv.) was quickly added. The flask was fitted with a three-way adapter connected to an $H_2$ balloon and a vacuum line. The system was purged three times with $H_2$ and then maintained under an $H_2$ atmosphere while stirring at ambient temperature for 36 h. The reaction mixture was filtered through Celite and concentrated *in vacuo,* affording **S-6** as a tan residue (74.2 mg, 78%). $^1H$ NMR (500 MHz, $CDCl_3$) δ: 5.41 (d, *J* = 9.6 Hz, 1H), 5.29 (d, *J* = 2.8 Hz, 1H), 5.20 (s, 1H), 5.15 (s, 1H), 5.12 – 5.02 (m, 1H), 4.88 (t, *J* = 9.9 Hz, 1H), 4.17 – 4.02 (m, 4H), 2.15 (t, *J* = 7.1 Hz, 6H), 2.05 (s, 2H), 1.98 (s, 2H), 1.32 (td, *J* = 7.0, 3.7 Hz, 9H).

**Synthesis of (2*R*,3*R*,4*S*,5*S*)-2-((*E*)-2-(diethoxyphosphoryl)vinyl)-6-(2,2,2-trichloro-1- imino-ethoxy)tetrahydro-2*H*-pyran-3,4,5-triyl triacetate (S-7).**

**S-6** (1.50 g, 3.40 mmol) was dissolved in DCM (15.0 mL) and added to an oven-dried flask flushed with $N_2$. Trichloroacetonitrile (3.40 mL, 34.0 mmol, 10 equiv) was added slowly, followed by 1,8-diazabicyclo[5.4.0]undec-7-ene (DBU, 0.600 mL, 0.340 mmol, 0.10 equiv) dropwise. Upon addition, the reaction mixture gradually turned deep brown. The reaction was deemed complete after 3 h of stirring at ambient temperature, as monitored by TLC (80% EtOAc/hexanes). The solvent was removed *in vacuo*, and the crude deep-brown oil was purified by silica gel flash chromatography (80% EtOAc/hexanes) to yield trichloroacetimidate **S-7** as a dark-brown, viscous oil (1.38 g, 70%). $^1$H NMR (400 MHz, $CDCl_3$) δ: 8.74 (s, 1H), 6.21 (d, *J* = 1.8 Hz, 1H), 5.44 (dt, *J* = 3.5, 1.8 Hz, 1H), 5.35 (ddd, *J* = 10.1, 3.5, 1.6 Hz, 1H), 5.20 (td, *J* = 10.0, 1.6 Hz, 1H), 4.10 – 4.03 (m, 4H), 3.94 (ddd, *J* = 10.0, 8.3, 2.6 Hz, 1H), 2.07 (s, 3H), 2.04 (s, 3H), 2.00 (s, 3H), 1.31 (td, *J* = 7.0, 1.5 Hz, 6H).$^{13}$C NMR (100 MHz, $CDCl_3$) δ: 170.34, 170.25, 170.16, 92.27, 77.39, 77.14, 76.89, 70.59, 69.66, 69.29, 68.47, 62.00, 60.53, 24.21, 21.50, 21.15, 21.06, 20.89, 20.83, 20.37, 16.50, 16.46, 14.27. HRMS (ESI-Pos) *m/z* calculated for $C_{19}H_{29}Cl_3O_{11}P$ (583.05): Found 606.0437 $[M+Na]^+$.

**Synthesis of (2-((2*R*,3*R*,4*S*,5*S*,6*S*)-3,4-diacetoxy-6-(prop-2-yn-1-yloxy)-5-((trimethylsilyl)-oxy)tetrahydro-2*H*-pyran-2-yl)ethyl)phosphonic acid (S-8).**

Trichloroacetimidate **S-7** (1.38 g, 3.40 mmol) was dissolved in DCM (20.0 mL) in an oven-dried flask and flushed with $N_2$. The reaction flask was cooled to 0 °C, and propargyl alcohol (0.980 mL, 17.0 mmol, 5.00 equiv) followed by trimethylsilyl trifluoromethanesulfonate (0.123 mL, 0.681 mmol, 0.200 equiv) were added dropwise. The reaction mixture was stirred vigorously on ice for 1.5 h and monitored by TLC (100% EtOAc). The reaction was quenched with saturated sodium bicarbonate, and the organic phase was dried over anhydrous sodium sulfate, filtered, and concentrated. The crude product was purified by silica gel flash chromatography (2–10% MeOH/DCM) to afford a viscous oil (0.323 g, 0.676 mmol).

The resulting oil was dissolved in anhydrous acetonitrile (8.00 mL), cooled to 0 °C, and treated with pyridine (0.600 mL, 7.43 mmol, 11.0 equiv), followed by bromotrimethylsilane (1.80 mL, 13.5 mmol, 20.0 equiv). The reaction was stirred at ambient temperature for 2 h, then quenched with pyridine (10.0 mL) and diluted with deionized $H_2O$. The mixture was concentrated with dioxane to afford a tan residue. The crude residue was dissolved in EtOAc and washed with brine (2 × 30.0 mL). The organic phase was dried over anhydrous sodium sulfate, filtered, and concentrated. The resulting oil was dissolved in acetic acid/milli-Q $H_2O$ (1:1 v/v, 8.00 mL), fitted with a condenser, and heated to 90 °C for 4 h with stirring. Reaction progress was monitored by TLC (5% MeOH/DCM). The solution was co-evaporated with Milli-Q $H_2O$ five times to remove excess acetic acid, then dried to afford propargyl glycoside **S-8** as a white solid (78.1 mg, 33%), which was carried forward to the next step without further purification. $^1$H NMR (500 MHz, $D_2O$) δ 5.33 – 5.25 (m, 1H), 5.20 – 5.09 (m, 2H), 5.09 – 4.92 (m, 2H), 4.47 – 4.26 (m, 2H), 4.19 – 4.00 (m, 2H), 2.92 (dp, $J$ = 6.7, 2.3 Hz, 1H), 2.19 (s, 2H), 2.22 – 2.13 (m, 2H), 2.13 – 2.04 (m, 1H), 2.01 (d, $J$ = 3.3 Hz, 1H), 1.97 – 1.88 (m, 2H), 1.92 (s, 2H), 1.79 – 1.67 (m, 2H), 1.25 (t, $J$ = 6.9 Hz, 1H). HRMS (ESI-Pos) *m/z* calculated for $C_{19}H_{29}Cl_3O_{11}P$ (583.05): Found 606.0437 $[M+Na]^+$.

**Synthesis of propargyl mannose 6-phosphonate glycoside (3).**

Crute propargyl glycoside **S-8** (0.0750 g, 0.118 mmol) was dissolved in MeOH (5.00 mL) in a scintillation vial. Sodium methoxide (0.0118 mmol) was added, and the reaction mixture was stirred at ambient temperature for 2 h. The reaction was quenched with DOWEX resin, filtered, and concentrated *in vacuo* to afford propargyl mannose-6-phosphonate **3** as a tan solid (52.7 mg, quant.). $^1$H NMR (500 MHz, $D_2O$) δ: 4.97 (d, $J$ = 1.8 Hz, 1H), 4.35 – 4.24 (m, 2H), 3.92 (dt, $J$ =

3.6, 1.8 Hz, 1H), 3.75 – 3.72 (m, 1H), 3.59 (td, $J$ = 9.1, 2.8 Hz, 1H), 3.50 (t, $J$ = 9.6 Hz, 1H), 2.90 (t, $J$ = 2.4 Hz, 1H), 2.15 – 2.05 (m, 1H), 2.03 – 1.93 (m, 1H), 1.83 – 1.71 (m, 2H). HRMS (ESI-Pos) $m/z$ calculated for $C_{19}H_{29}Cl_3O_{11}P$ (583.05): Found 606.0437 $[M+Na]^+$.

#### Synthesis of glycopolymers *p*(Man6Pn)-biotin and *p*(Man)-biotin.

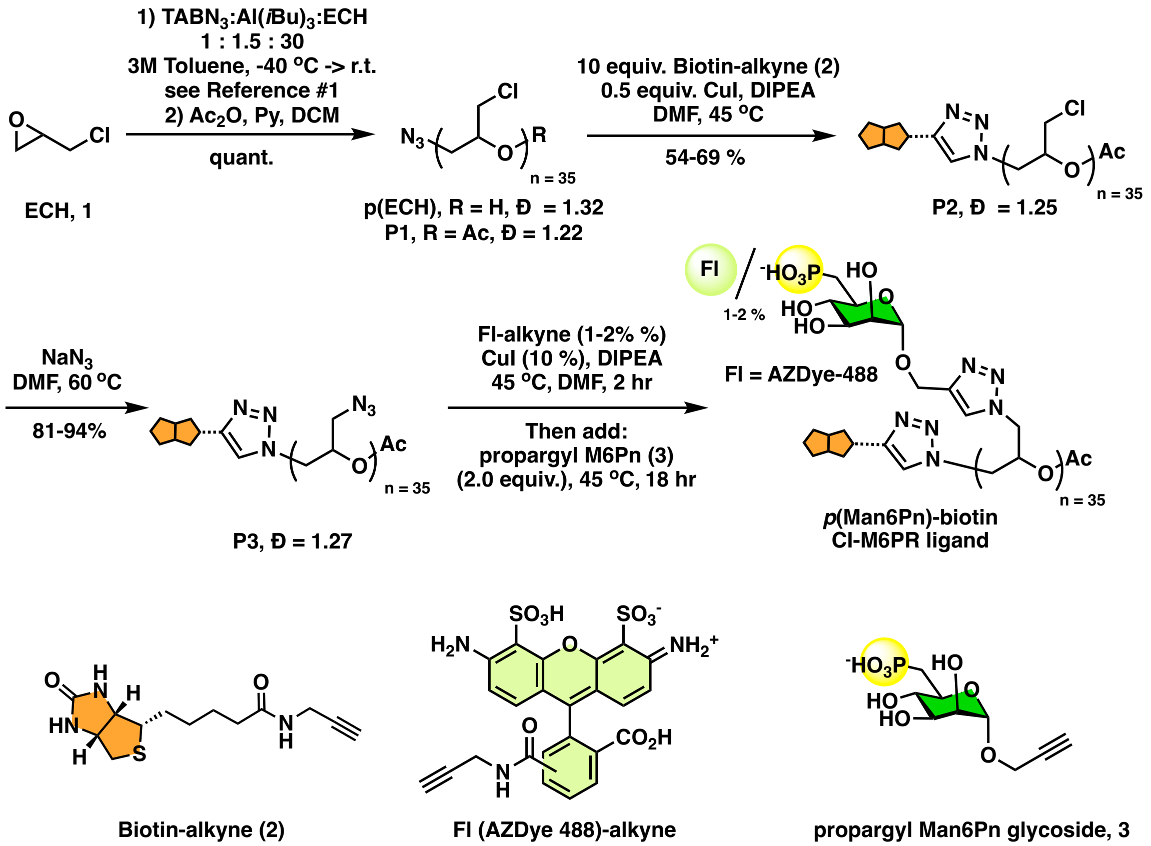


**Scheme S2.** Synthesis of glycopolymers ***p*(Man6Pn)-biotin** and ***p*(Man)-biotin**.

##### Synthesis of chain-end *O*-acetylated poly(epichlorohydrin) backbone P1.

In a flame-dried Schlenk flask (10.0 mL), ***p*(ECH)** polymer precursor[2] (Mn = 3056 g·mol$^{-1}$, PD = 35, Đ = 1.32; 79.3 mg, 0.0259 mmol) was dissolved in degassed anhydrous DCM (0.79 mL). Pyridine (0.10 mL, 50.0 equiv) was added, followed by $Ac_2O$ (0.12 mL, 50.0 equiv). The reaction was stirred at room temperature for 25 h. The reaction mixture was evaporated and dried under vacuum to afford polymer **P1** as a clear oil (81.2 mg, quantitative yield). $^1H$ NMR (500 MHz,

$CDCl_3$) δ: 3.77 – 3.56 (m, 208H), 2.13 – 2.03 (m, 3H). SEC: 0.7 mL/min in DMF (0.2% LiBr, 70 °C): Mn = 3224, Đ = 1.22, DP = 35.

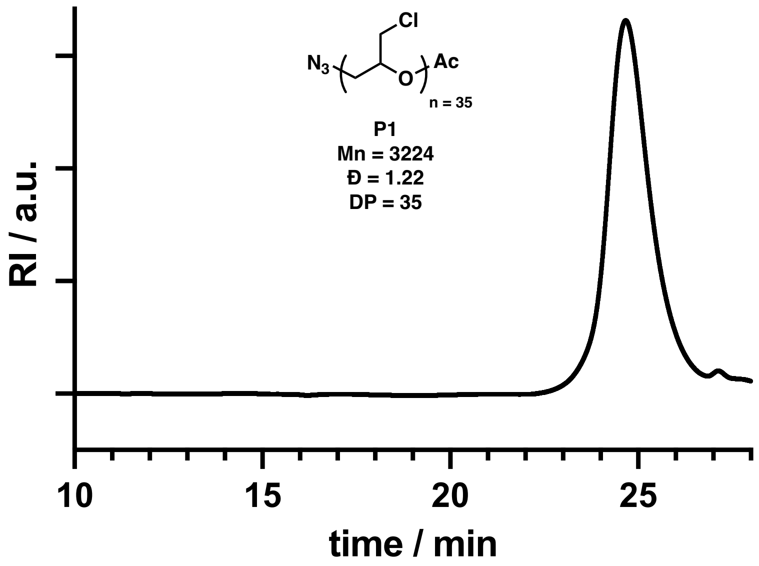


**Figure S1.** GPC trace of chain-end *O*-acetylated poly(epichlorohydrin) backbone **P1**.

**Synthesis of chain-end-biotinylated poly(epichlorohydrin) polymer P2.**

Polymer **P1** (53.3 mg, 0.0174 mmol) was dissolved in degassed anhydrous DMSO (0.2 mL). Biotin propargyl amide (48.9 mg, 0.174 mmol, 10.0 equiv) was added, followed by CuI (1.70 mg, 0.00870 mmol, 0.50 equiv) and DIPEA (3 μL, 0.00870 mmol, 1.0 equiv). The reaction mixture was stirred in the dark at 40 °C for 22 h. Degassed anhydrous DMSO (0.3 mL) was then added, and the reaction mixture was stirred in the dark at 40 °C for an additional 26.0 h. The mixture was cooled to room temperature, and Cuprisorb beads (~1.00 g) were added. The resulting suspension was stirred at room temperature for 19.0 h. The solution was filtered through Celite to remove the beads and washed with DCM. After brief evaporation to remove DCM, $H_2O$ was added to the polymer solution to precipitate the polymer. The precipitate was dissolved in a small amount of acetone, and $H_2O$ was added again to reprecipitate the polymer. The resultant polymer was dried under vacuum to yield **P2** as a viscous oil (44.3 mg, 83.0% yield). $^1H$ NMR (400 MHz, $CDCl_3$) δ:

7.71 (s, 1H), 3.80 – 3.55 (m, 200H). SEC: 0.7 mL/min in DMF (0.2% LiBr, 70 °C): Mn = 3222, Đ = 1.24, DP = 35.

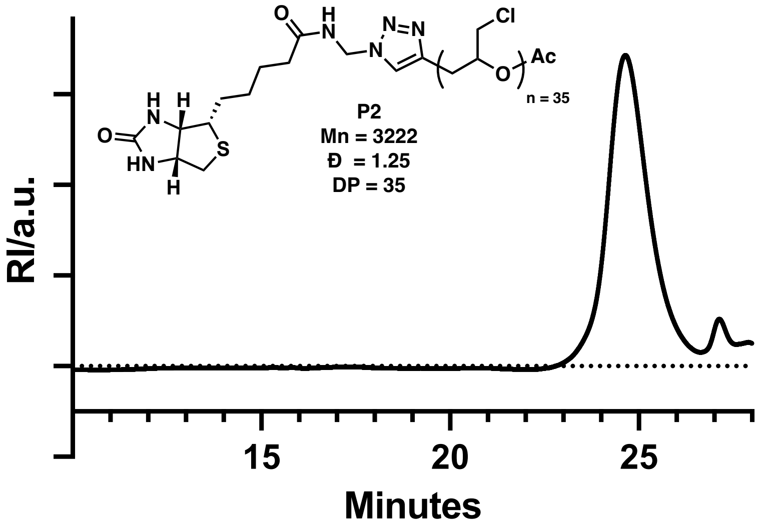


**Figure S2.** GPC trace of chain-end biotinylated poly(epichlorohydrin) polymer **P2**.

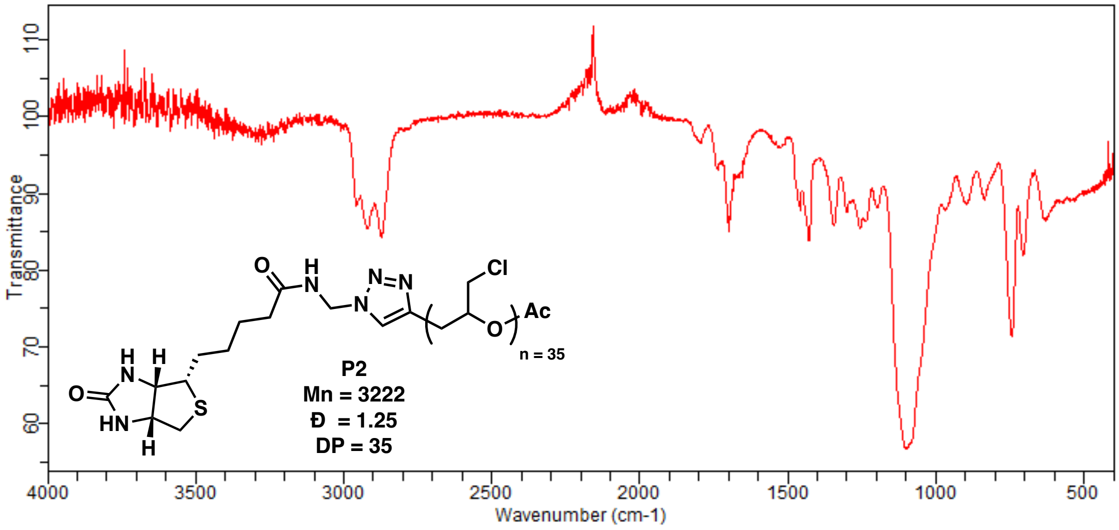


**Figure S3.** IR spectrum of chain-end biotinylated poly(epichlorohydrin) polymer **P2**.

**Synthesis of chain-end biotinylated poly(glycidyl azide) polymer P3.**

In a flame-dried Schlenk flask (10.0 mL), polymer **P2** (41.7 mg, 0.451 mmol) was dissolved in anhydrous DMF (0.3 mL). To this solution was added $NaN_3$ (58.6 mg, 0.901 mmol, 2.0 equiv). The reaction mixture was stirred at 80 °C for 22.5 h under argon to ensure complete conversion.

The mixture was cooled to room temperature and filtered through Celite. The polymer solution was precipitated in acetone and $H_2O$, yielding **P3** as a clear, viscous oil (41.9 mg, 94.0% yield). $^1$H NMR (400 MHz, $CDCl_3$) δ: 3.80 – 3.55 (m, 150H), 3.53 – 3.32 (m, 50H), 2.12 (d, $J$ = 1.1 Hz, 3H). SEC: 0.7 mL/min in DMF (0.2% LiBr, 70 °C): Mn = 3749, Đ = 1.27, DP = 35.

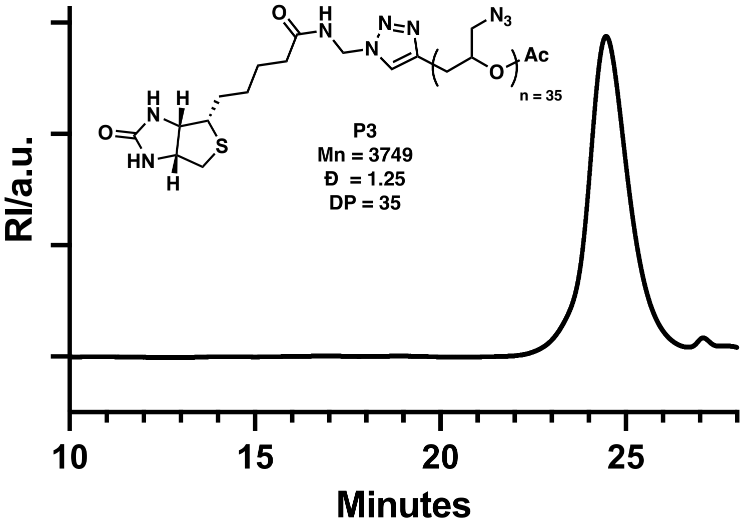


**Figure S4.** GPC trace of chain-end biotinylated poly(glycidyl azide) polymer **P3**.

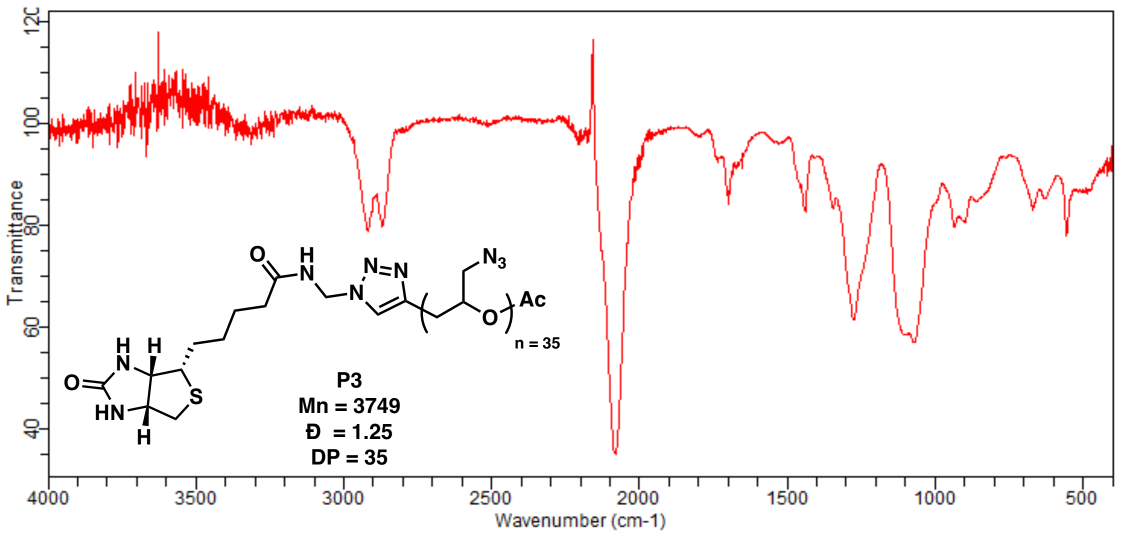


**Figure S5.** IR spectrum of chain-end biotinylated poly(glycidyl azide) polymer **P3**.

**Synthesis of CI-M6PR glycopolymer ligand *p*(M6Pn)-biotin.**

Poly(glycidyl azide)-biotin **P3** (1.82 mg, 0.0184 mmol) was dissolved in degassed anhydrous DMSO (0.1 mL). To this solution was added AZDye-488 alkyne (0.310 mg, 0.551 μmol, 0.03 equiv per side chain) in degassed anhydrous DMSO (31 μL), followed by CuI (0.350 mg, 1.84 μmol, 0.10 equiv per side chain) and DIPEA (3.20 μL, 0.0184 mmol, 1.00 equiv per side chain) in degassed anhydrous DMSO (88 μL). The reaction mixture was stirred in the dark at 40 °C for 25 h. After this time, propargyl-M6Pn glycoside (3, 10.8 mg, 0.0364 mmol, 2.00 equiv) in degassed anhydrous DMSO (0.15 mL) was added, followed by CuI (0.350 mg, 1.84 μmol, 0.10 equiv) and DIPEA (3.20 μL, 0.0184 mmol, 1.00 equiv) in degassed anhydrous DMSO (88 μL). The reaction mixture was stirred in the dark at 60 °C for 23 h. The solution was cooled to room temperature, and Cuprisorb beads (~1.00 g) together with DI water (~1.00 mL) were added. The mixture was stirred at room temperature overnight. The solution was filtered through Celite to remove the beads. After size-exclusion purification with a PD-10 column, the polymer solution was lyophilized. The dry residue was triturated three times with methanol. The resulting polymer was dried under vacuum to yield the AZDye-488–labeled ***p*(M6Pn)-biotin** (7.60 mg, quantitative yield; 1.3% dye incorporation per monomer via UV-Vis absorbance at $\lambda_{max} = 488$). Glycoside attachment was estimated to be quantitative on the complete disappearance of the characteristic azide stretch at 2100 $cm^{-1}$ in IR, and the theoretical Mn of the final *p*(M6Pn)-biotin polymer was calculated accordingly.

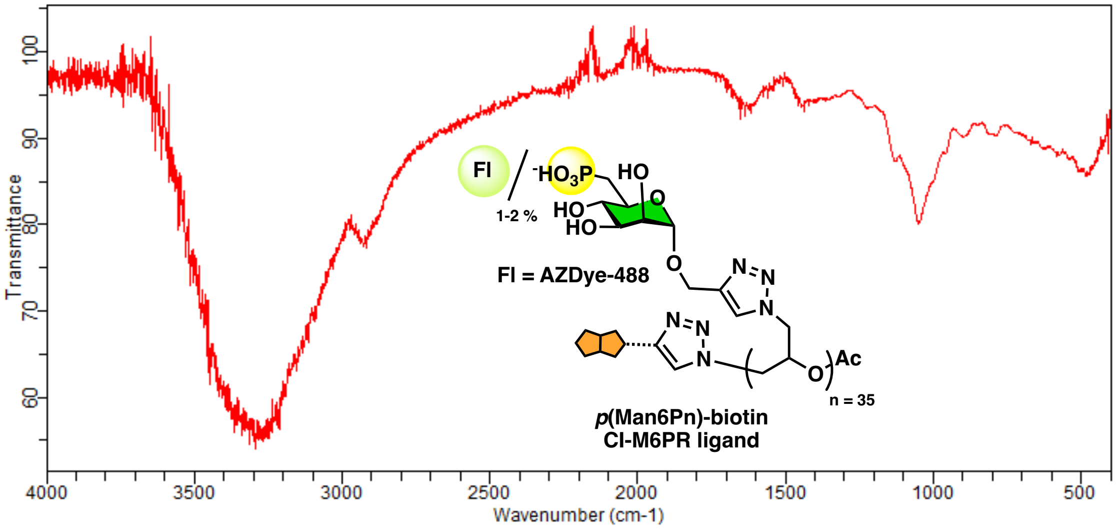


**Figure S6.** IR spectrum of CI-M6PR glycopolymer ligand ***p*(M6Pn)-biotin**.

**Synthesis of CI-M6PR non-binding glycopolymer *p*(Man)-biotin.**

Poly(glycidyl azide)-biotin **P3** (2.78 mg, 0.0280 mmol) was dissolved in degassed anhydrous DMSO (0.10 mL). To this solution was added AZDye-488 alkyne (0.480 mg, 0.842 μmol, 0.03 equiv per side chain) in degassed anhydrous DMSO (48.0 μL), followed by CuI (0.500 mg, 2.63 μmol, 0.09 equiv per side chain) and DIPEA (4.90 μL, 0.0280 mmol, 1.00 equiv per side chain) in degassed anhydrous DMSO (50.0 μL). The reaction mixture was stirred in the dark at 40 °C for 23 h. After this time, α-propargyl mannoside[4] (9.20 mg, 0.042 mmol, 1.50 equiv per side chain) was added to the solution, followed by CuI (0.500 mg, 2.63 μmol, 0.09 equiv per side chain) and DIPEA (4.90 μL, 0.0280 mmol, 1.00 equiv per side chain) in degassed anhydrous DMSO (50.0 μL). The reaction mixture was stirred in the dark at 40 °C for 27.0 h. The solution was cooled to room temperature, and Cuprisorb beads (~1.00 g) together with DI water (~1.50 mL) were added. The mixture was stirred at room temperature overnight. The solution was filtered through Celite to remove the beads. After purification with a NAP-25 column, the polymer solution was lyophilized. The dry residue was triturated three times with methanol. The resulting polymer was dried under vacuum to yield the AZDye-488–labeled ***p*(Man)-biotin** (7.39 mg, 83.0% yield; 2.2%

dye incorporation per monomer via UV-Vis absorbance at $\lambda_{max}$ = 488). Glycoside attachment was estimated to be quantitative based on the complete disappearance of the characteristic azide stretch at 2100 cm$^{-1}$ in IR, and the theoretical Mn of the final ***p*(Man)-biotin** polymer was calculated accordingly.

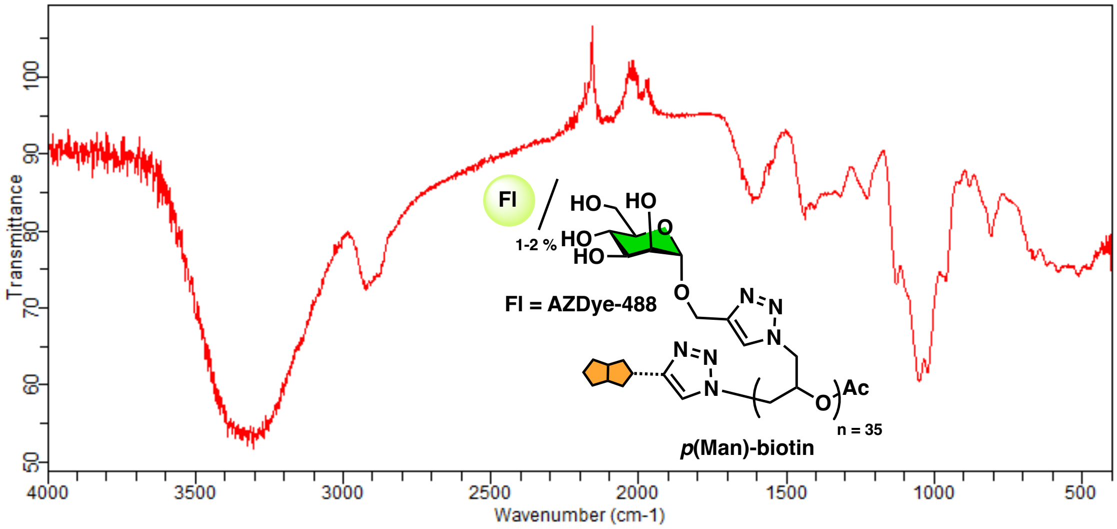


**Figure S7.** IR spectrum of CI-M6PR non-binding glycopolymer ***p*(Man)-biotin**.

#### GLYTAC assembly and characterization

##### Reducing end-biotinylation of heparin (Hep-biotin)

Synthesis of **Hep-biotin** by reductive amination of was carried out as previously described.[5] In a PCR tube, heparin sodium salt (Iduron, 6.96 mg, 0.464 µmol, 1.00 equiv) was dissolved in sodium acetate buffer (570 µL, 100 mM sodium acetate, pH 5.0). A solution of diaminopropane in sodium acetate buffer (1.00 M, 9.28 µL, 9.28 µmol, 20.0 equiv) was added, followed by a solution of sodium cyanoborohydride in sodium acetate buffer (32.0 mM, 116 µL, 3.71 µmol, 8.00 equiv). The reaction was heated at 37 °C for 24 h. After this time, the mixture was purified on a PD-10 column following the manufacturer's protocol (Cytiva) to remove unreacted linker, concentrated by spin filtration against water (3k MWCO, 12 000 × g, 8.00 min, 6 rounds), and lyophilized to afford the amine end-functionalized heparin (**Hep-NH$_2$**) product as a white solid (6.96 mg, quantitative yield).

Next, the **Hep-NH$_2$** (6.96 mg, 0.464 μmol) was dissolved in sodium phosphate buffer (0.400 mL, 100 mM $Na_3PO_4$, 150 mM NaCl, pH 8.0) in a PCR tube, and NHS-PEG$_4$-biotin was added (5.50 mg, 9.28 μmol, 20.0 equiv). The reaction proceeded at ambient temperature for 24 h, followed by purification on a PD-10 column according to the manufacturer's protocol (Cytiva) to remove unreacted linker. The product was concentrated by spin filtration against water (3k MWCO, 12 000 × g, 8.00 min, 6 rounds) and lyophilized to afford the reducing end-biotinylated heparin (**Hep-biotin**) as a white solid (4.45 mg, 63.9% yield).

The extent of heparin chain biotinylation in **Hep-biotin** was determined using a HABA assay (Pierce Biotin Quantification Kit). A solution of HABA–avidin premix was reconstituted in ultrapure water, and the absorbance at 500 nm was recorded. After addition of a known amount of **Hep-biotin** (20.0 μM final concentration, based on dry weight of the desalted polysaccharide), the change in absorbance was measured and used to calculate the molar concentration of biotin in the sample. The fraction of heparin chains end-functionalized with biotin was determined as the ratio of biotin to heparin concentrations: ~14.5 %.

**Assembly of *p*(M6Pn)-SA 1:1 complex.**

Streptavidin (**SA**, Leinco, 2.00 mg/mL, 500 μL) was mixed with ***p*(M6Pn)-biotin** (533 μL, 144 μM, 4.0 equiv) in DPBS buffer, pH 7.4. The reaction mixture was rotated in the dark for 24 h. The resulting mixture was purified by spin filtration (4.00 mL spin filter, 50 kDa MWCO, 4000 × g) for 12 cycles, washing with DPBS+/+. The obtained conjugate was stored in PBS (410 μL) and analyzed by gel electrophoresis.

For gel electrophoresis, agarose gels were cast at 1.50% (w/v) using low-EEO biology grade agarose (Millipore) dissolved in 1×Tris–borate EDTA (Life Technologies). Gels were cast in an EasyCast OWL electrophoresis chamber and run with ice-cold 1× Tris–borate EDTA (Life Technologies) at 180 V for 45 min. Samples were loaded into wells in 1:2 glycerol in ultrapure water to obtain a final glycerol concentration of 10.0% (v/v). AF488 fluorescence was detected using an Amersham A680 RGB imager, and ImageJ was used to analyze collected images. UV–Vis absorbance at 280 nm and 494 nm was measured using a NanoDrop One to determine **SA** and ***p*(M6Pn)-biotin** concentrations, respectively, thereby establishing the 1:1 ***p*(M6Pn)-SA** complex stoichiometry.

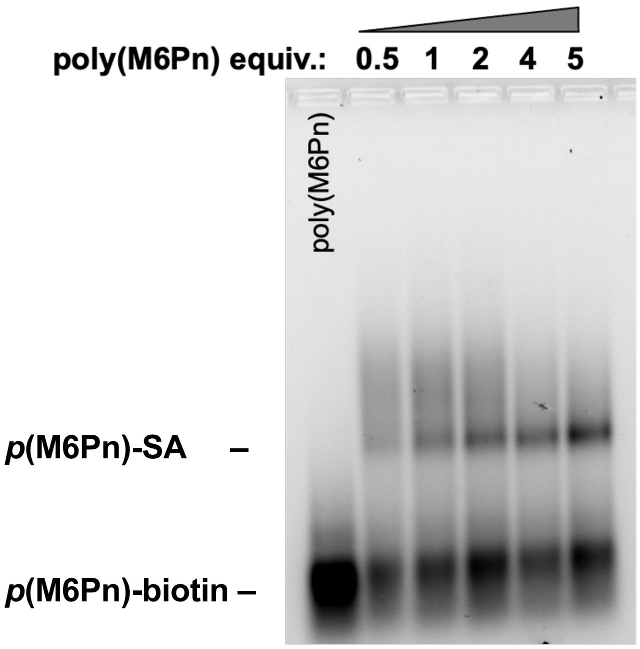


**Figure S8.** Optimization of ***p*(M6Pn)-SA** complex assembly. Agarose gel electrophoresis (1.5% agarose w/v in 1X TBE, 180V, 60 min) was used to monitor ***p*(M6Pn)-SA** formation. ***p*(M6Pn)-biotin** was incubated with **SA** at increasing ratios (0.5 – 5.0 equiv.). Formation of conjugate resulted in an upward mobility shift based on ***p*(M6Pn)-biotin** fluorescence (AF488).

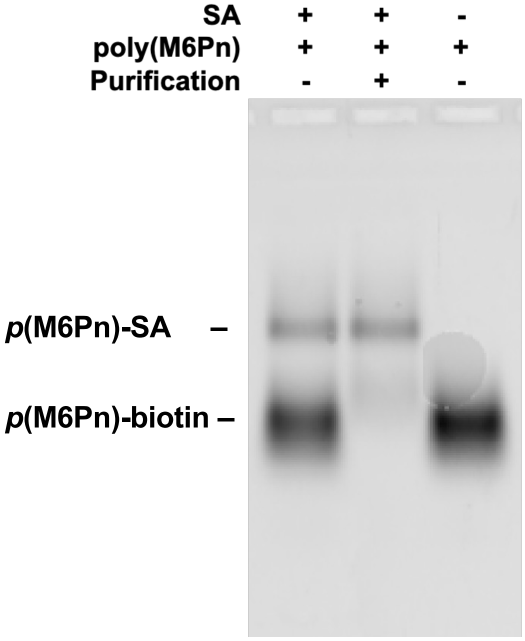


**Figure S9.** Analysis of purified ***p*(M6Pn)-SA** conjugate. Agarose gel electrophoresis (1.5% agarose w/v in 1x TBE, 180V, 45 min) was used to determine the formation and purity of the ***p*(M6Pn)-SA** complex after spin filtration. Due to molecular weight addition, **SA** complexation to ***p*(M6Pn)-biotin** resulted in a modest upward shift based on ***p*(M6Pn)-biotin** fluorescence (AF488).

**Assembly of Hep-SA-*p*(Man6Pn) GLYTAC conjugate**

***p*(M6Pn)–SA** (410 µL, 11 µM) was mixed with **Hep-biotin** (311 µL, 29 µM based on 14.5% chain biotinylation, 2.6 equiv). The reaction mixture was incubated in the dark for 24 h. The resulting mixture was purified by spin filtration (4.0 mL spin filter, 50 kDa MWCO, 4000 × g) for 12 cycles, washing with DPBS+/+. The purified **Hep-SA-*p*(Man6Pn)** conjugate was stored in PBS (80 µL) and analyzed by gel electrophoresis (1.50% agarose, 180 V, 45.0 min), as described above. The gel was then visualized by AF488 fluorescence. AF488 fluorescence was used to determine the concentration of **p(M6Pn)-biotin,** and **Hep-biotin** to **SA** stoichiometry was characterized by disaccharide analysis followed by (GRIL) LTQ–MS.

**Determination of hep/SA complex stoichiometry via (GRIL) LTQ–MS disaccharide analysis**

A **Hep–SA**-solution (20.0 µM **SA**, based on UV–Vis) was treated with Heparinase I, II, and III at 37 °C for 18 h. Following digestion, proteins were removed using 10 kDa MWCO filters, and the HS disaccharides were dried. The HS disaccharides were dissolved in aniline and reacted with sodium cyanoborohydride in DMSO:acetic acid (7:3, v/v). The reaction was carried out at 65 °C for 1 h, followed by 16 h at 37 °C. The sample was then dried and analyzed by glycan reductive isotope labeling (GRIL) LTQ–MS to determine the amount of disaccharide in the preparation (1.53 µg). The average molecular weight of heparin (15.0 kDa, per manufacturer) was used to determine Hep concentration (20.4 µM Hep) in the analyzed sample (5 µL), giving a final **hep–biotin**/**SA** ratio of 1.02.

| | | | | | | | |
|---|---|---|---|---|---|---|---|
| | | | Amount Loaded (µL) | | | | |
| | added to vial= | 5 | total in vial= | 10 | amt injected= | 4 | |
| | | | Injected | In Digest | | In Digest | |
| | 12C Ion Intensity | 13C Ion Intensity | pmole | pmole | Mole % | ng | |
| D0H0 | 1.47 | 19.2 | 0.31 | 3.06 | 0.07 | 1.03 | |
| D0A0 | 120 | 31.3 | 15.34 | 153.35 | 3.39 | 58.12 | |
| D0H6 | 6.6 | 42.3 | 0.62 | 6.24 | 0.14 | 2.60 | |
| D2H0 | 3.05 | 35.4 | 0.34 | 3.45 | 0.08 | 1.44 | |
| D0S0 | 202 | 48.4 | 16.69 | 166.94 | 3.69 | 69.61 | |
| D0A6 | 243 | 35.8 | 27.15 | 271.51 | 6.00 | 124.62 | |
| D2A0 | 60.1 | 34.3 | 7.01 | 70.09 | 1.55 | 32.17 | |
| D2H6 | 26.2 | 51.7 | 2.03 | 20.27 | 0.45 | 10.07 | |
| D0S6 | 467 | 31.3 | 59.68 | 596.81 | 13.18 | 296.61 | |
| D2S0 | 388 | 53.5 | 29.01 | 290.09 | 6.41 | 144.18 | |
| D2A6 | 111.4 | 61.81 | 7.21 | 72.09 | 1.59 | 38.86 | |
| D2S6 | 1425.3 | 19.837 | 287.40 | 2874.02 | 63.47 | 1658.31 | |
| total | | | | 4527.93 | 100.00 | 1525.91 | |
| | | | | | Total in Digest = | 1.53 | µg |
| | | | | | Total in Prep = | 1.53 | µg |
| | [Hep] = 1.53 µg / (15000 µg/µmol) / 5µL | | | | | | |
| | [Hep] = 0.0000204 M | | | | % N, 2-O, 6-O-sulfated | | |
| | [Hep] = 20.4 µM | | | | N-SO3 | 2-O-SO3 | 6-O-SO3 |
| | Hep:SA = 20.4 / 20 | | | | 86.75 | 73.54 | 84.83 |
| | Hep:SA = 1.02 | | | | | | |
| | | | | | 2.45 | average SO3 per disaccharid | |

**Table S1.** Disaccharide composition of **Hep-SA** conjugate.

#### BIOLOGICAL PROCEDURES

##### Cell Culture

Hela cells were cultured following standard tissue culture practices using reagents and supplements purchased from ThremoFisher Scientific or as indicated in the methods sections and used as received according to manufacturer's recommendation. HeLa cells were maintained in Dulbecco's Modified Eagle Medium (DMEM) supplemented with 10% Fetal Bovine Serum (FBS) and 1% Penicllin/Stroptemycin.

##### Internalization and lysosomal targeting of *p*(M6Pn) in HeLa cells by microscopy.

HeLa cells were seeded at 0.4 × 106 cells/well in a 35 mm dish (Mattek P35G-1.5-10-C) in growth media. After 24 h, cells were washed with DPBS+/+ and then incubated in growth medium supplemented with GLYTAC (5 μM) for 30 minutes at 37˚C. To each dish, LysoTracker Deep Red (Invitrogen L12492) was added, and incubation continued for an additional 30 min at 37˚C. Following incubation, cells were washed three times with DPBS+/+ and stained with DAPI for 10 min at room temperature, then washed three more times. Finally, cells were fixed with 4% PFA for 8 min and washed three times with DPBS, and stored in 100 mM HEPES/DPBS buffer for fluorescence imaging.

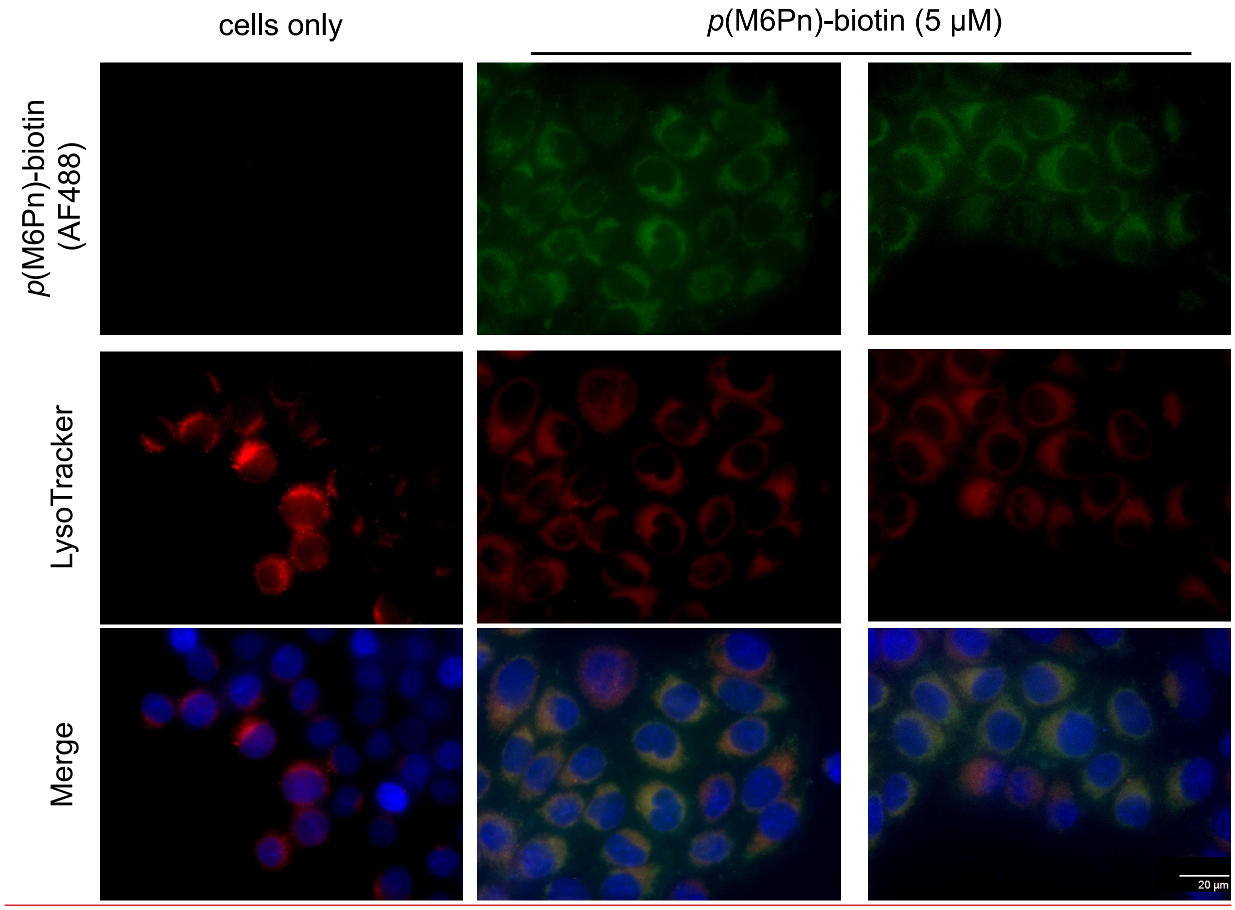


**Figure S10.** Internalization of ***p*(M6Pn)** (5.0 μM) in HeLa cells via microscopy (Extended data for **Figure 2B**).

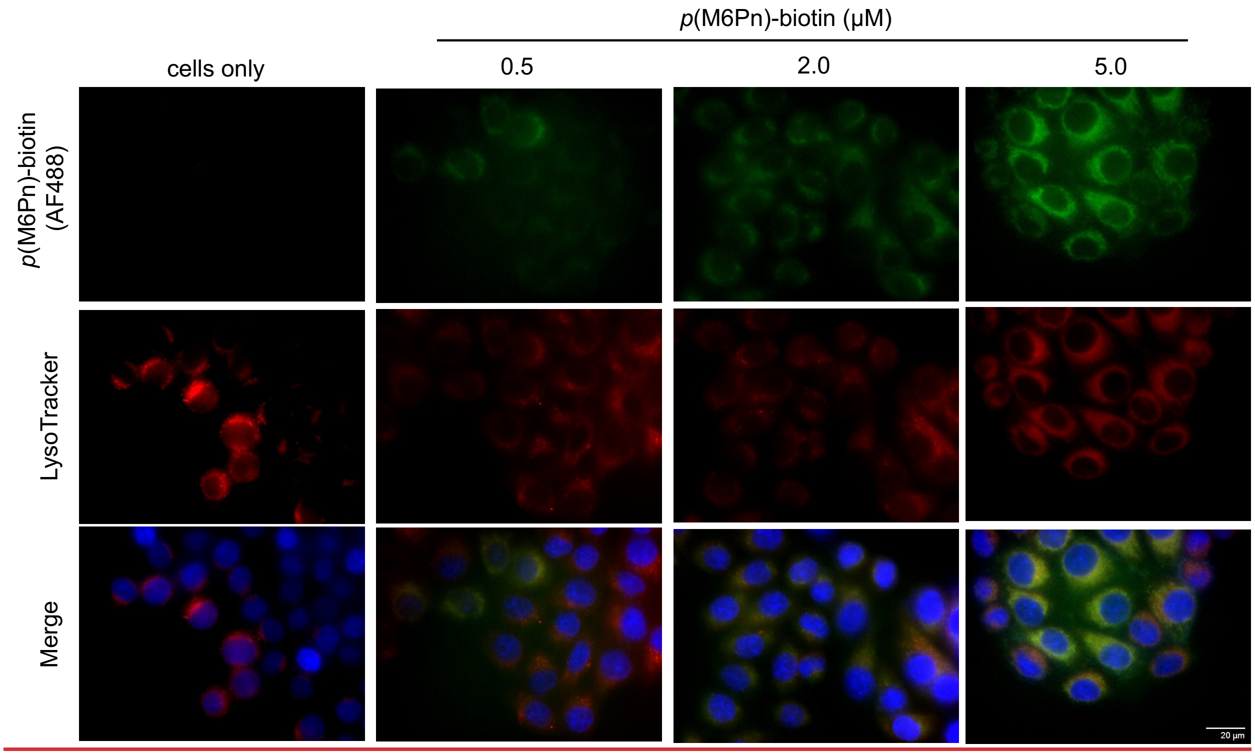


**Figure S11.** Dose-dependent internalization of ***p*(M6Pn)** (0.5, 2.0, and 5.0 μM) in HeLa cells via microscopy. HeLa cells treated with increasing concentrations of ***p*(M6Pn)-biotin** (0.5-5.0 μM, 1 hr, 37 °C) exhibited dose-dependent colocalization with the LysoTracker dye by fluorescence microscopy.

#### *p*(M6Pn)-mediated SA647 uptake by flow cytometry.

HeLa cells were seeded at 0.25 x $10^5$ cells/$cm^2$ in a 24-well plate. After 48 h, the media was aspirated, and the cells, at ~70% confluence, were washed once with DPBS$^{+/+,}$ then added growth medium supplemented with SA-647 (0.5–5 μM) or SA-647 with glycopolymer (0.5–5 μM), and immediately transferred to 37 °C with 5% $CO_{2.}$ After 1 h, the medium was aspirated, the cells were washed once with DPBS$^{+/+,}$ and the cells were lifted with trypsin-EDTA (0.05%). The cells were then transferred to a 96-well V-bottom plate. Cells were washed 3 times with 0.5% BSA/D-PBS$^{+}$ and analyzed by flow cytometry.

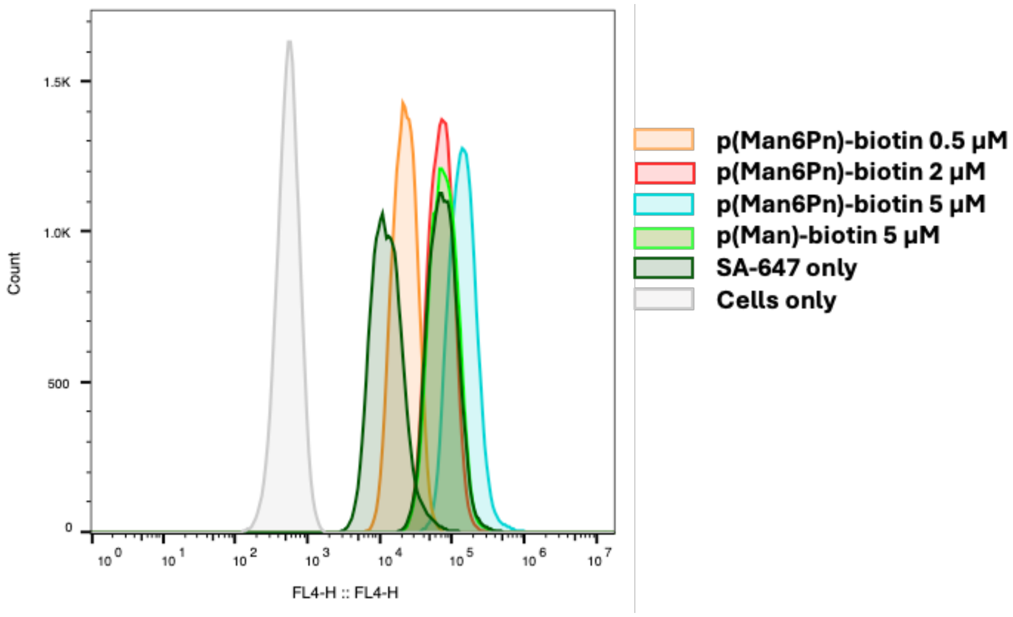


**Figure S12.** Uptake of SA-647 in HeLa cells mediated by equimolar ***p*(M6Pn)-biotin**. Flow cytometry analysis of HeLa cells treated with a 1:1 mixture of **SA-647** and ***p*(M6Pn)-biotin** at increasing concentration (0.0 – 5.0 μM) for 1 hr at 37˚C.

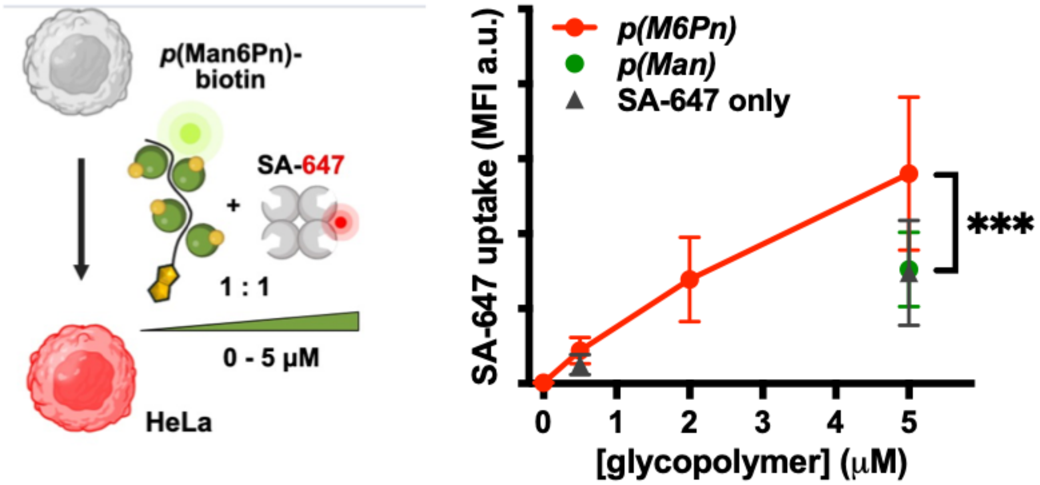


**Figure S13.** Uptake of SA-647 in HeLa cells mediated by equimolar ***p*(M6Pn)-biotin.** Flow cytometry analysis of HeLa cells treated 1:1 mixture of SA-647 and ***p*(M6Pn)-biotin** at increasing concentration (0.5, 2.0, and 5.0 μM) for 1 h at 37˚C.

#### FGF2 degradation immunohistochemistry (IHC).

HeLa cells were seeded at 0.33 × $10^{\wedge 5}$ cells/well in an 8-well slide (ibidi u-Slide 8 well) in growth media. After 48 h, cells were washed with DPBS$^{+/+}$ and then incubated in growth medium supplemented with FGF2 (200 nM) and GLYTAC (2 μM) or Hep-SA (2 μM) for the specified duration. Following incubation, cells were washed 3 times with DPBS, then fixed with 4% PFA for 8 min and washed 3 times with DPBS. Cells were permeabilized with 0.3% Tween-20 in 0.1% Saponin in 100 mM HEPES/DPBS for 15 min. Cells were washed thrice and blocked in 3% BSA in 0.1% Saponin in 100 mM HEPES/DPBS for 30 min at ambient temperature. Following three washes with DPBS, mouse anti-FGF2 (1:500) in 0.1% BSA in 0.1% Saponin in 100 mM HEPES in DPBS was bound overnight at 4˚C on an orbital shaker. Cells were washed 3 times and incubated with anti-mouse AF647 (1:1000) in 0.1% Saponin in 100 mM HEPES/DPBS for 1 h at ambient temperature. Following three washes, anti-LAMP2-AF555 (1:500) in in 0.1% Saponin in 100 mM HEPES/DPBS was incubated for 1 h at ambient temperature. Finally, cells were stained with DAPI for 10 min at ambient temperature, washed thrice, and stored in 100 mM HEPES/DPBS buffer for fluorescence imaging.

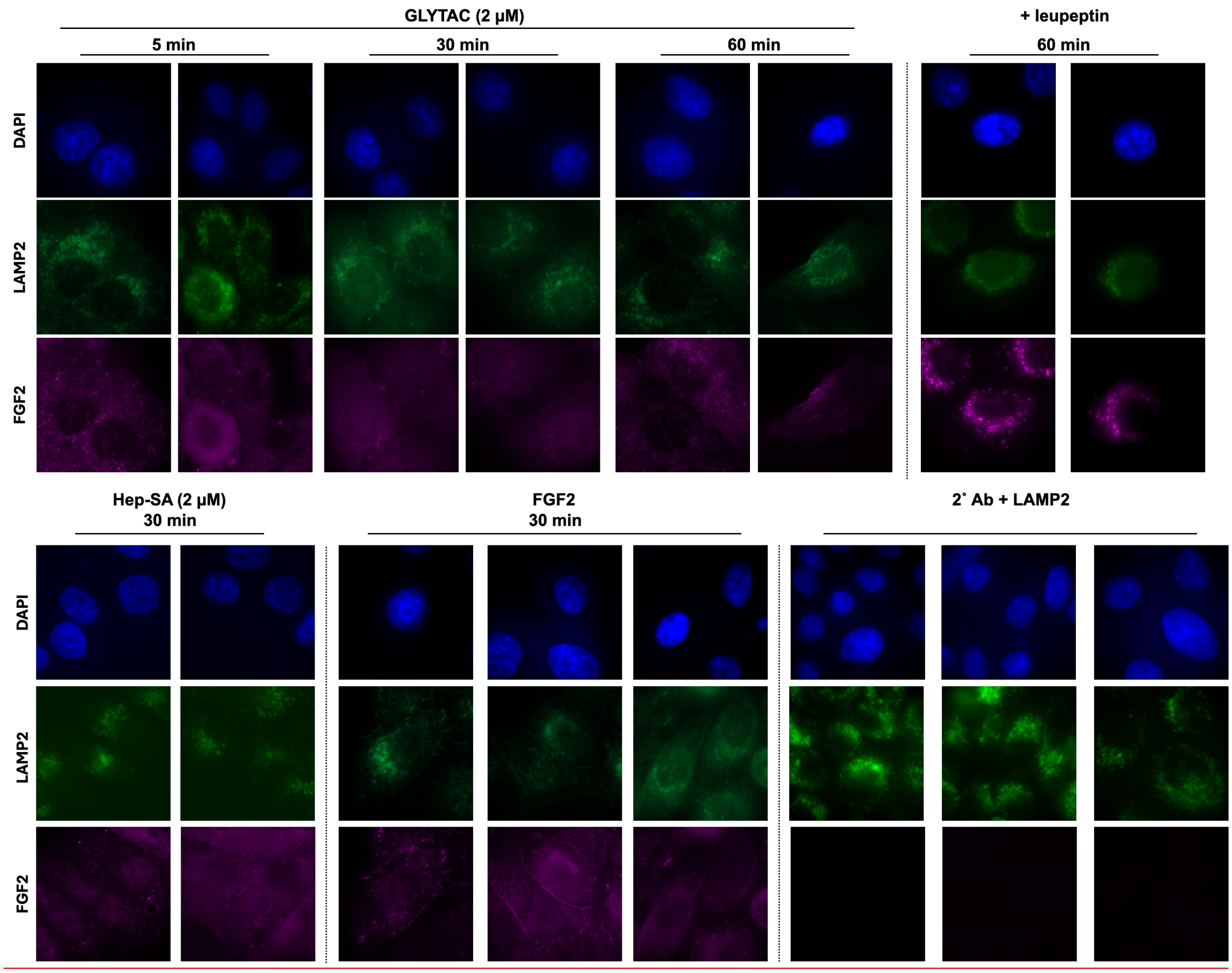


**Figure S14.** Internalization of FGF2 in HeLa cells (expanded data for **Figure 4B**). Fluorescent confocal images of HeLa immunostained for FGF2 and lysosome via LAMP2. Cells were treated with **GLYTAC** (2 µM) or **Hep-SA** (2 µM) with FGF2 (200 nM) over a period of 60 min.

#### FGFR degradation by Western Blotting

HeLa cells were seeded at 0.1 × $10^6$ cells/mL in 0.5 mL per well of a 24-well plate in DMEM + 10% FBS + 1% P/S. After 24 h, the growth media was replaced with 0.5 mL DMEM for 18 hours of serum starvation. Media was removed, cells were washed once with DPBS, and then replaced with DMEM + Hep/GLYTAC (0–2000 nM) containing FGF2 (20 ng/mL) for the indicated time period. Following incubation, cells were washed once with DPBS then lysed for 5 min in lysis buffer (1x RIPA, 1x PMSF, 1x protease inhibitor cocktail). Cell lysate was sonicated briefly and then clarified by centrifugation at 14,000 × g for 10 min at 4 °C. Lysate was analyzed for protein content using the BCA assay. 10 μg of protein was separated on a 10% SDS-PAGE gel and transferred to a PVDF membrane for Western blotting. The membrane was blocked in TBST containing 5% BSA for 1 hr at room temperature. Following 3 washes with TBST, primary antibodies (1:1000 in 5% BSA/TBST) were incubated overnight at 4 °C. The membrane was then washed 3 times with TBST and incubated with HRP-conjugated secondary antibodies (1:8000 in 5% BSA/TBST) for 1 h at room temperature. Membranes were visualized with Luminata Forte HRP detection reagent on Amersham A680 RGB image.

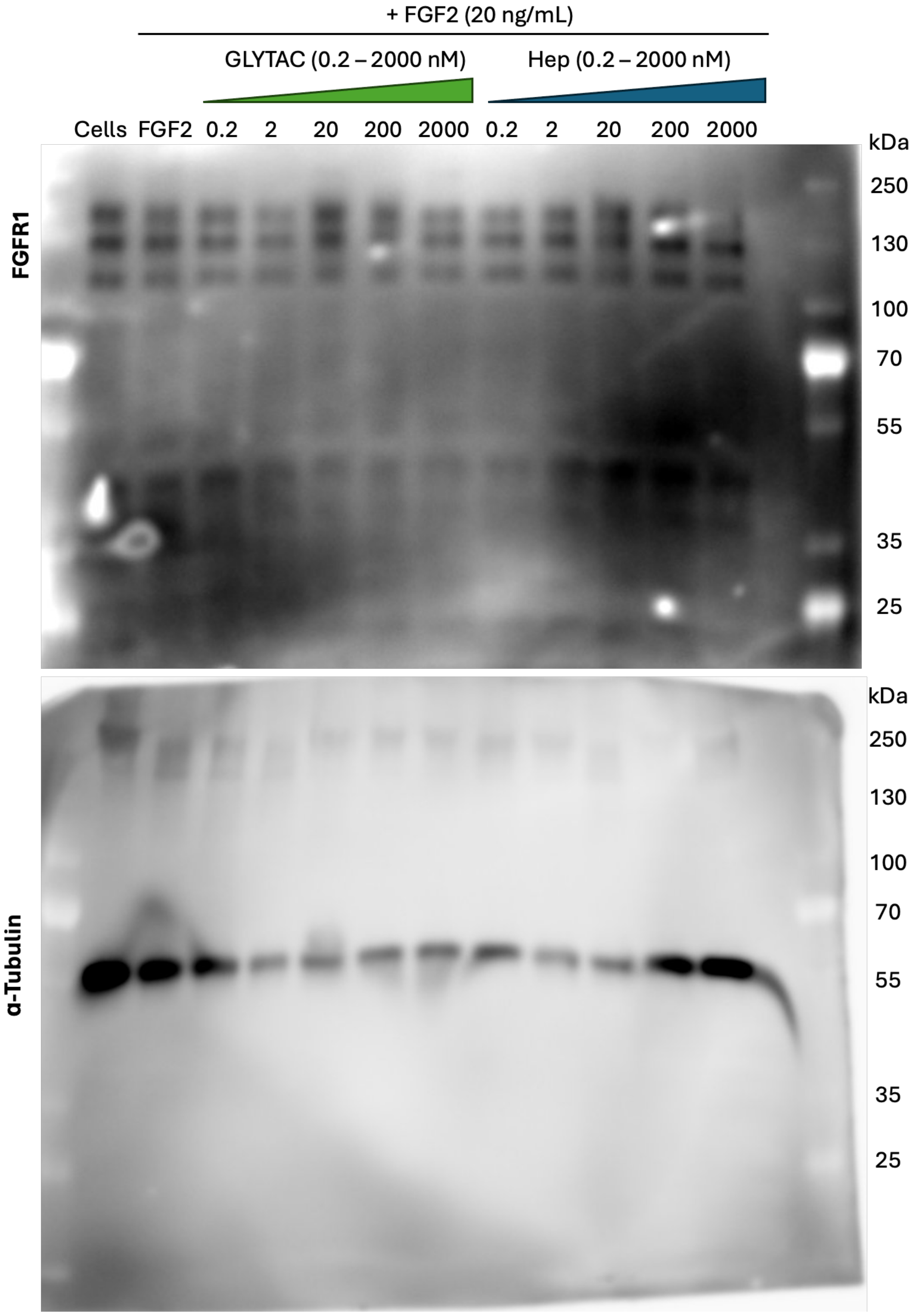


**Figure S15.** Western blot analysis of FGFR1 levels in HeLa cells. FGFR1 and ∝-Tubulin levels were assessed in HeLa cells treated Hep-SA or GLYTAC (0.2 – 2000 nM) in the presence of FGF2 (20 ng/mL).

### APPENDIX: $^{1}$H and $^{13}$C NMR spectra of synthetic compounds

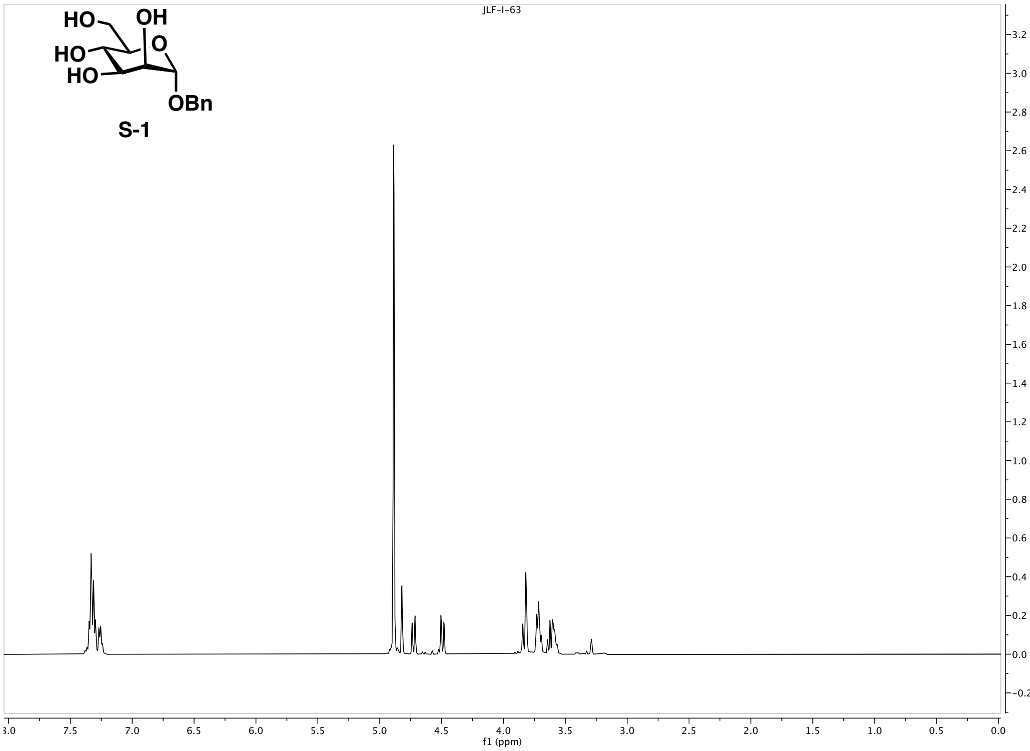


**Figure S-A.1.** $^{1}$H NMR (500 MHz, $CD_3OD$) spectrum for compound **S-1**.

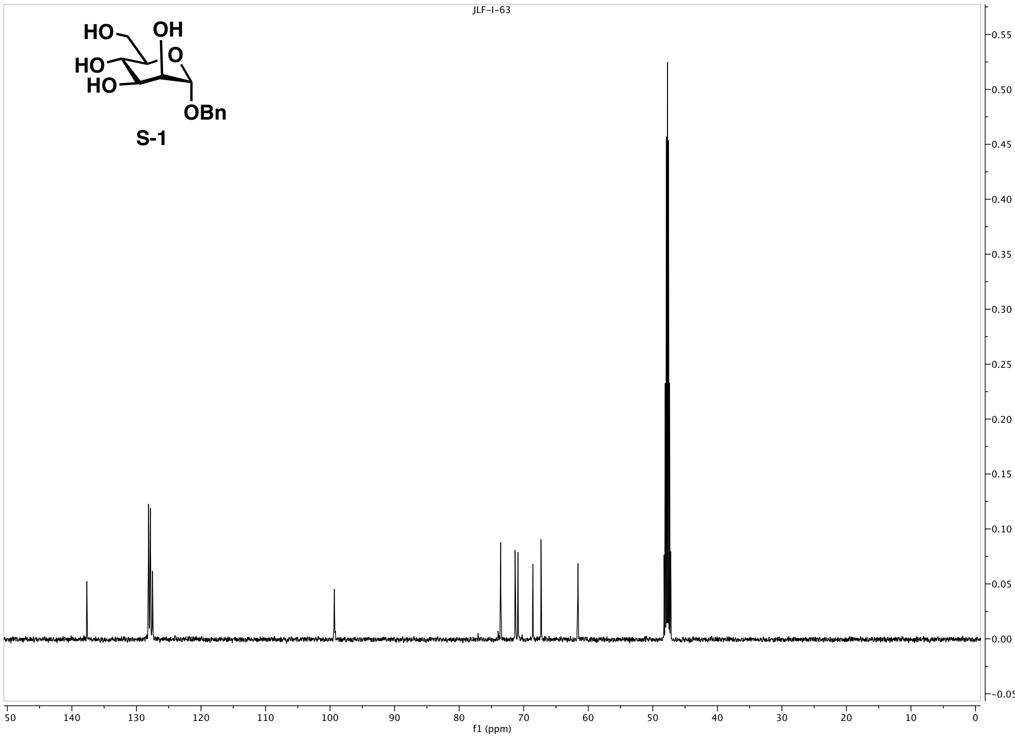


**Figure S-A.2** $^{13}$C NMR (126 MHz, $CD_3OD$) spectrum for compound **S-1**.

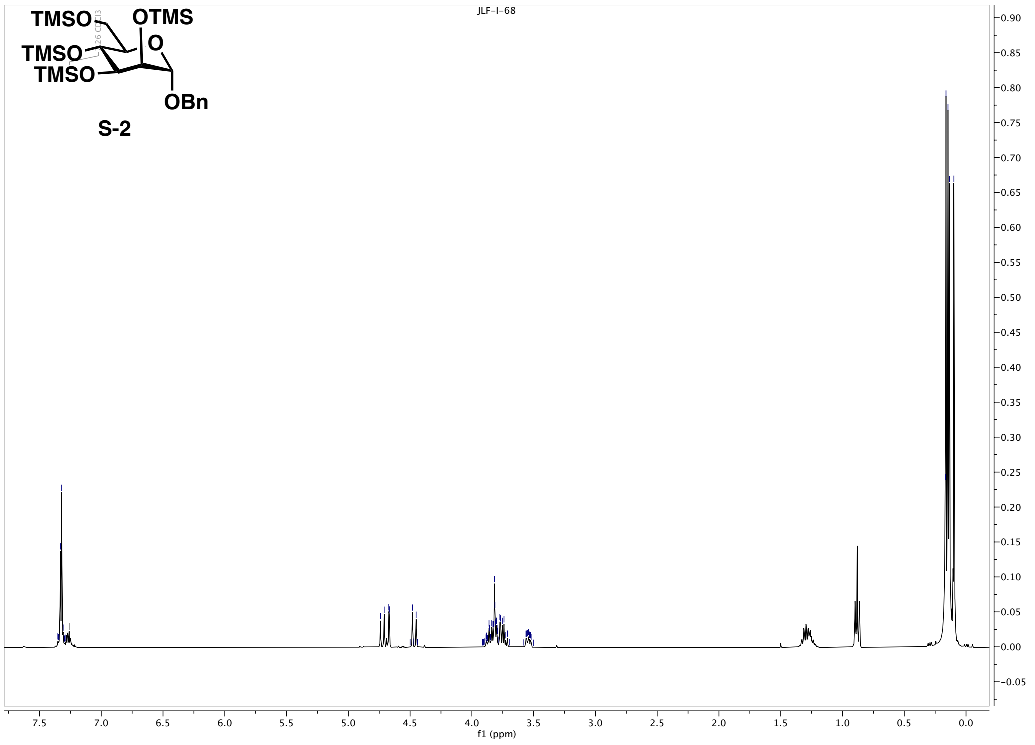


**Figure S-A.3.** $^{1}H$ NMR (500 MHz, $CDCl_3$) spectrum for compound **S-2**.

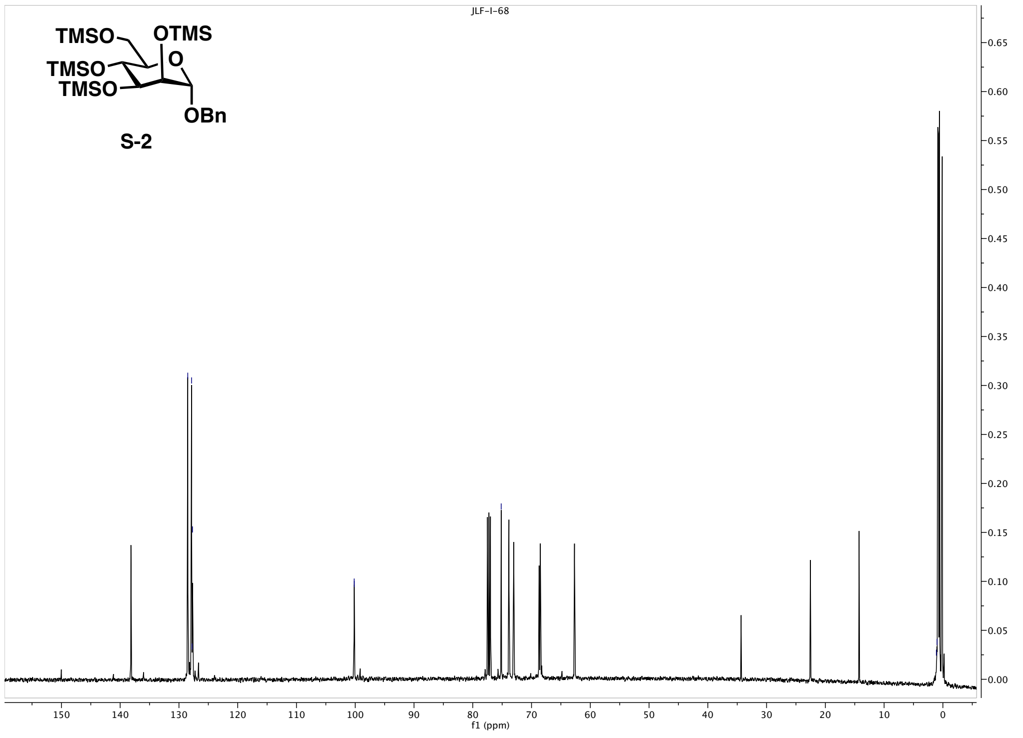


**Figure S-A.4.** $^{13}C$ NMR (126 MHz, $CDCl_3$) spectrum for compound **S-2**.

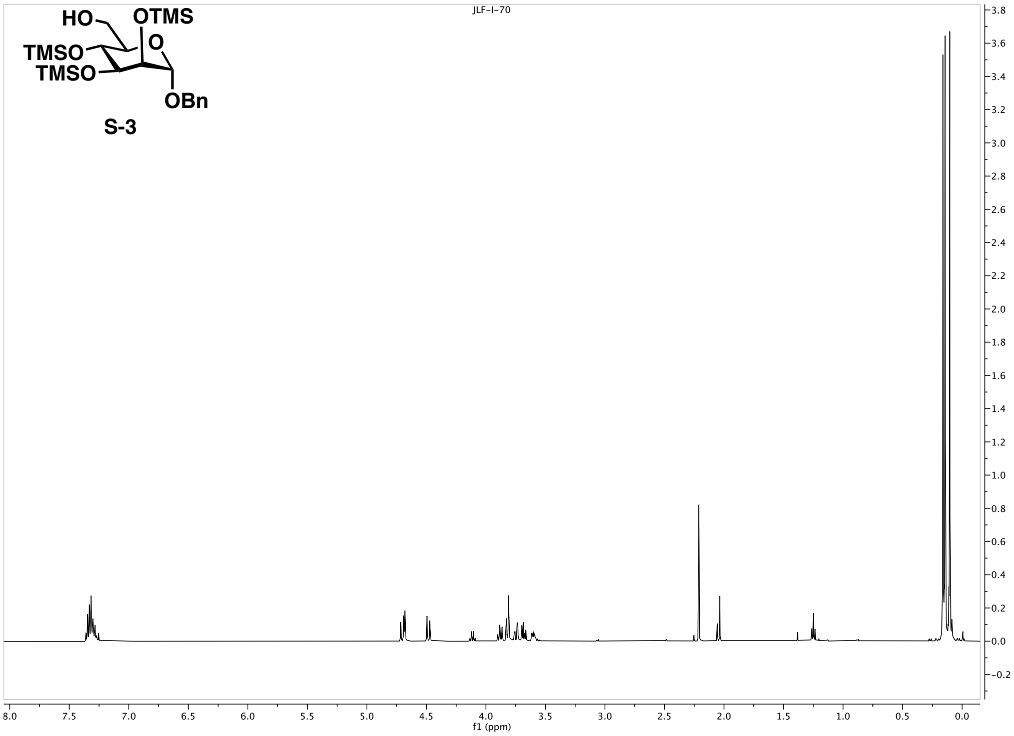


**Figure S-A.5.** $^{1}$H NMR (500 MHz, $CDCl_3$) spectrum for compound **S-3**.

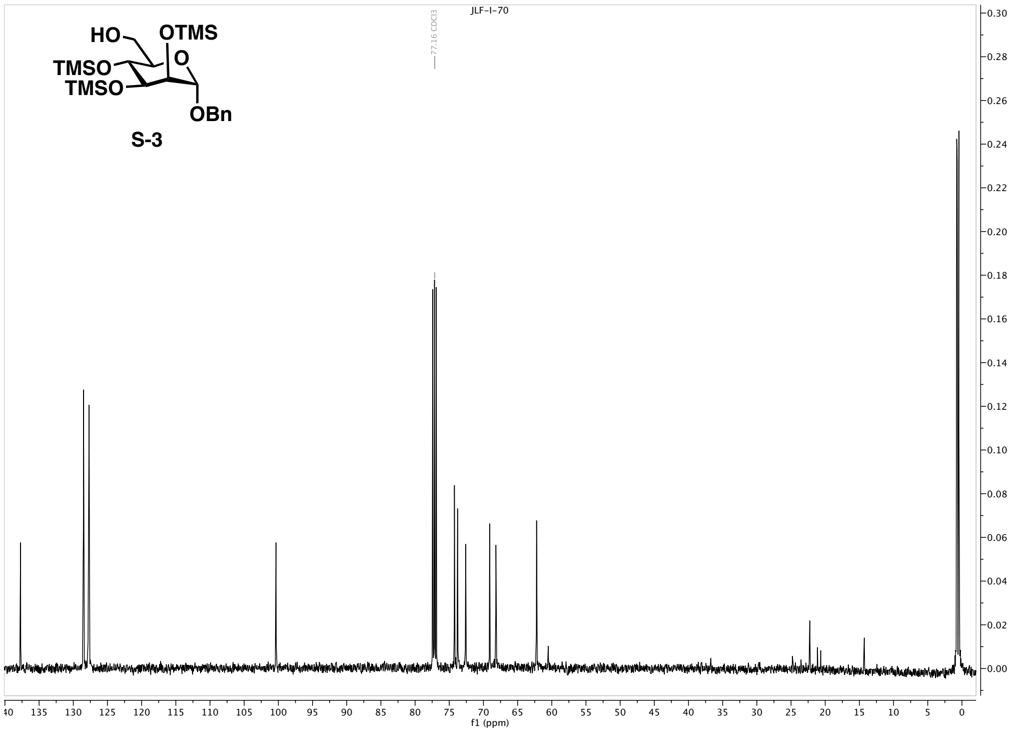


**Figure S-A.6.** $^{13}$C NMR (126 MHz, $CDCl_3$) spectrum for compound **S-3**.

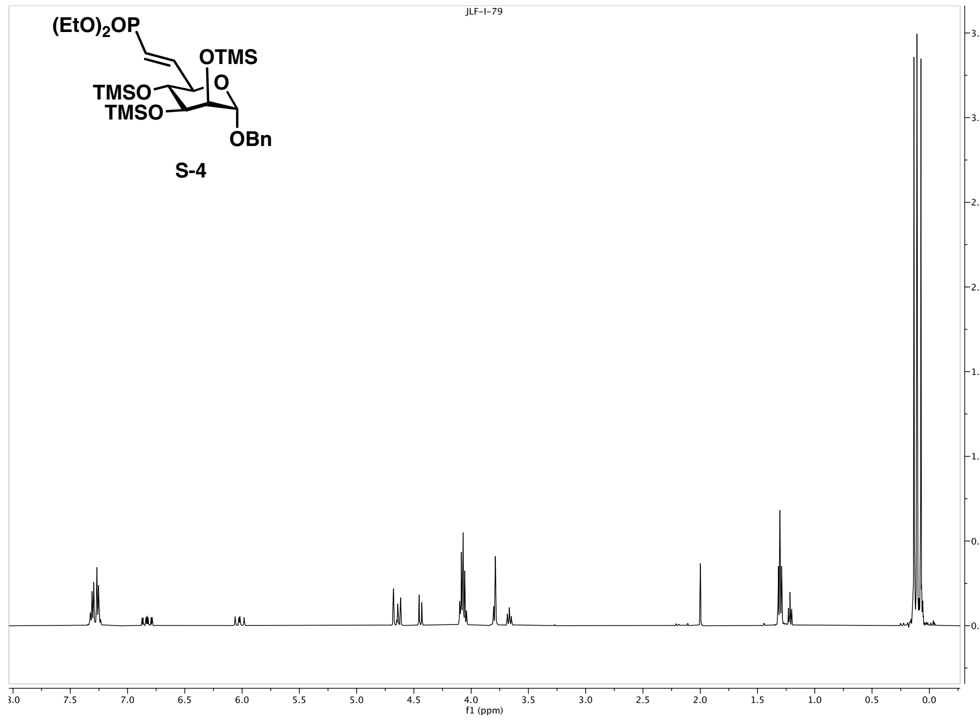


**Figure S-A.7.** $^1$H NMR (500 MHz, $CDCl_3$) spectrum for compound **S-4**.

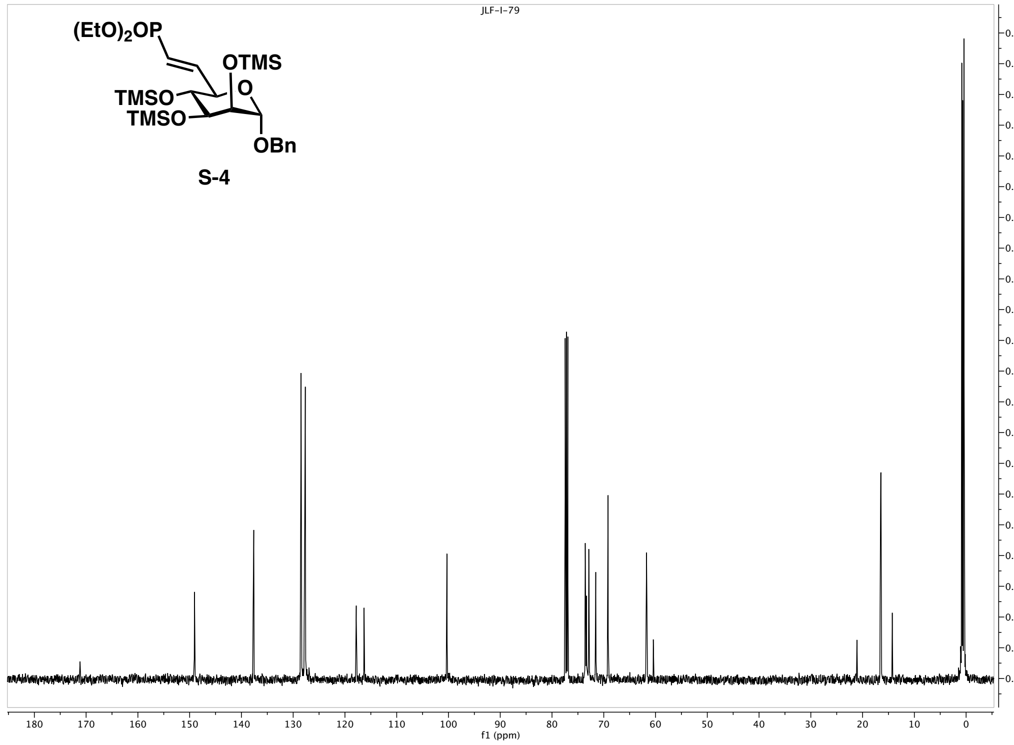


**Figure S-A.8.** $^{13}$C NMR (126 MHz, $CDCl_3$) spectrum for compound **S-4**.

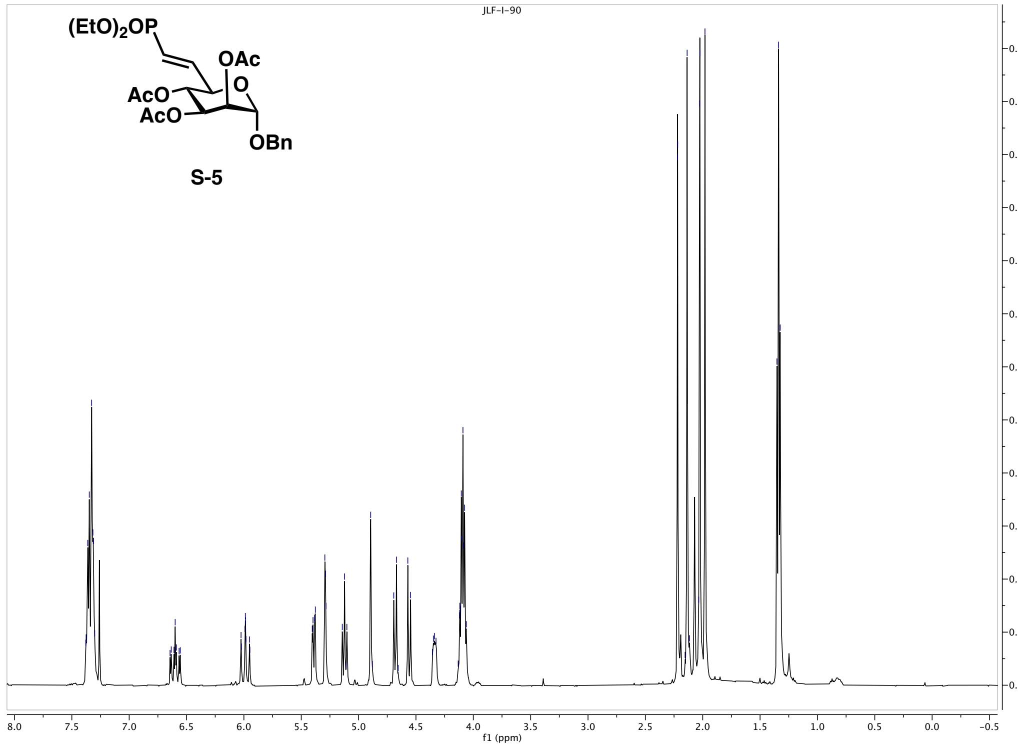


**Figure S-A.9.** $^{1}$H NMR (500 MHz, $CDCl_3$) spectrum for compound **S-5**.

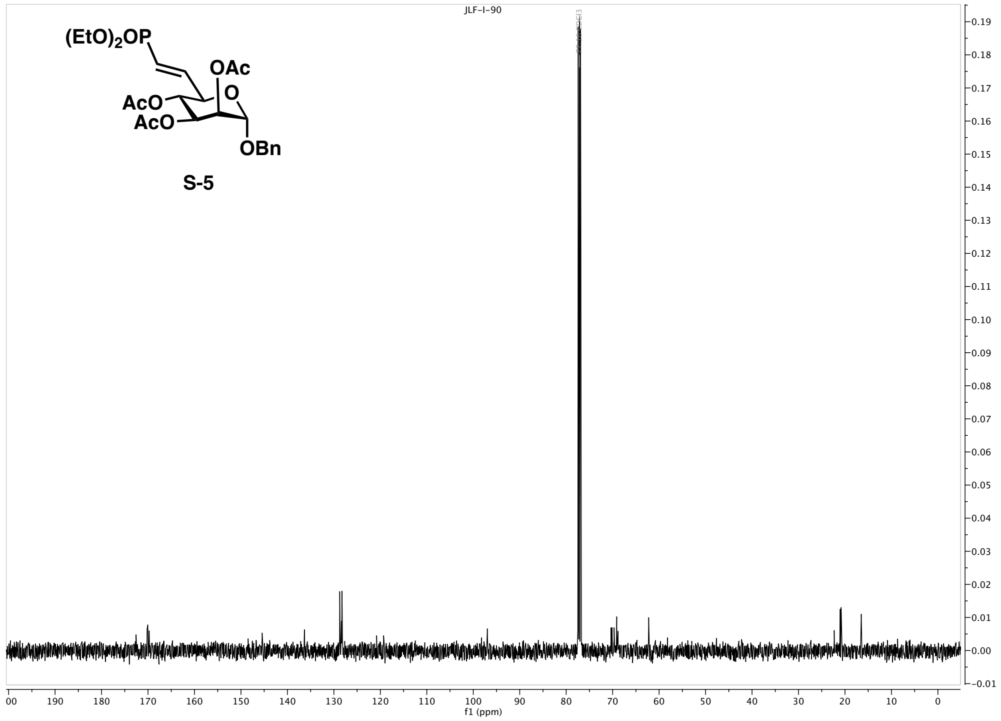


**Figure S-A.10.** $^{13}$C NMR (500 MHz, $CDCl_3$) spectrum for compound **S-5**.

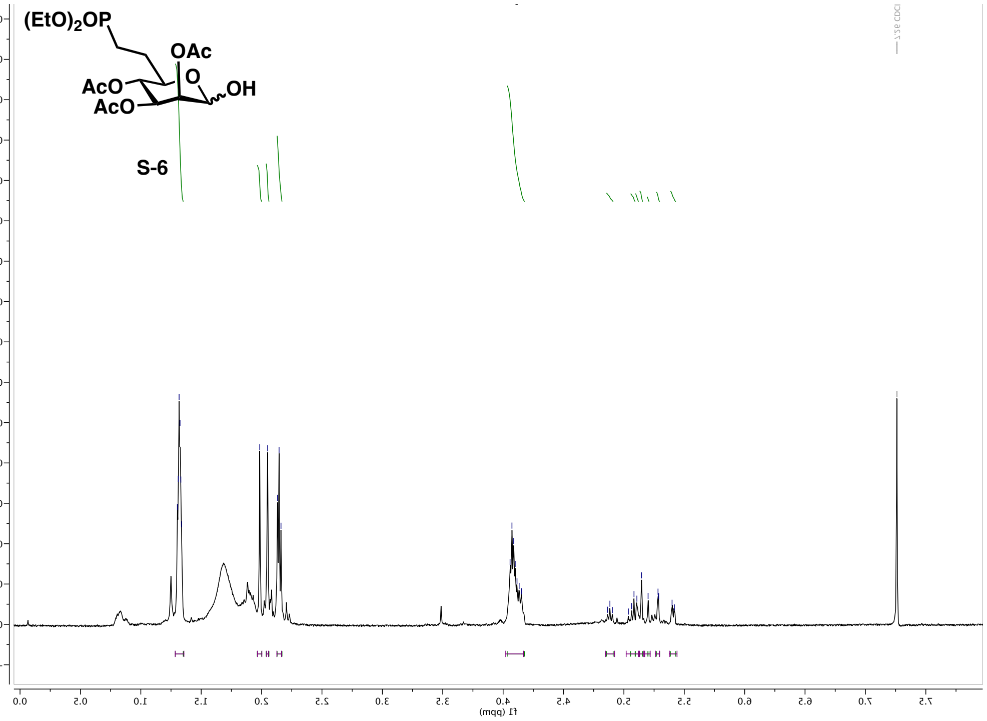


**Figure S-A.11.** $^1$H NMR (500 MHz, CDCl$_3$) spectrum for compound **S-6**.

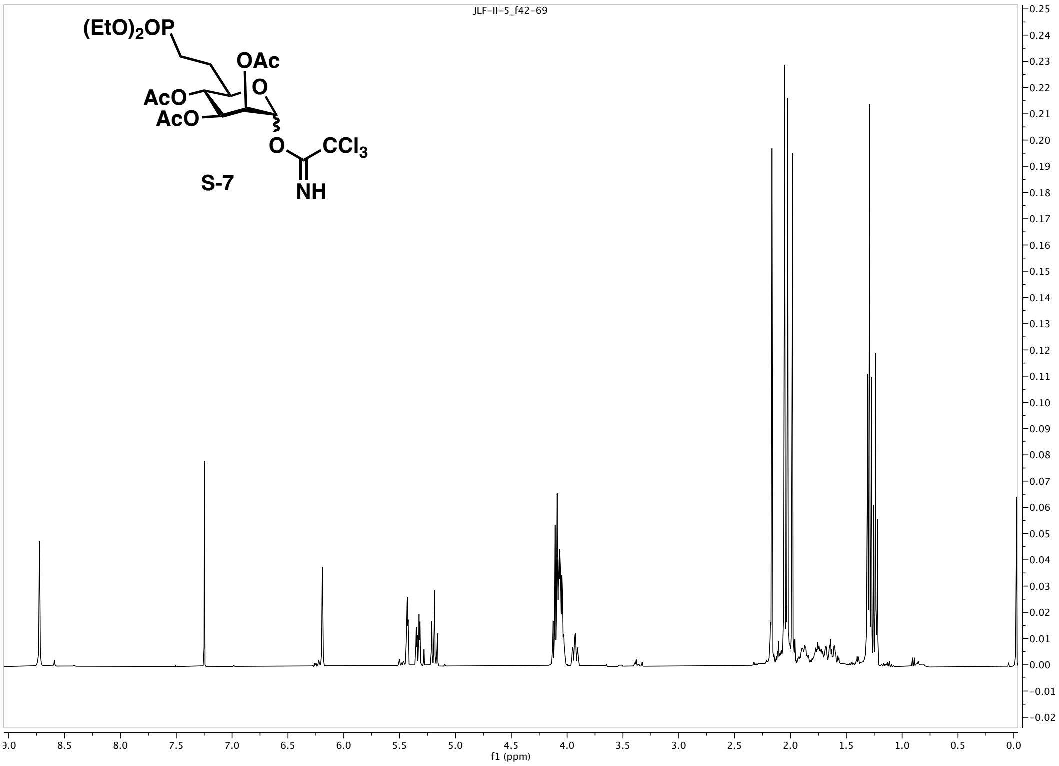


**Figure S-A.12.** $^1$H NMR (400 MHz, CDCl$_3$) spectrum for compound **S-7**.

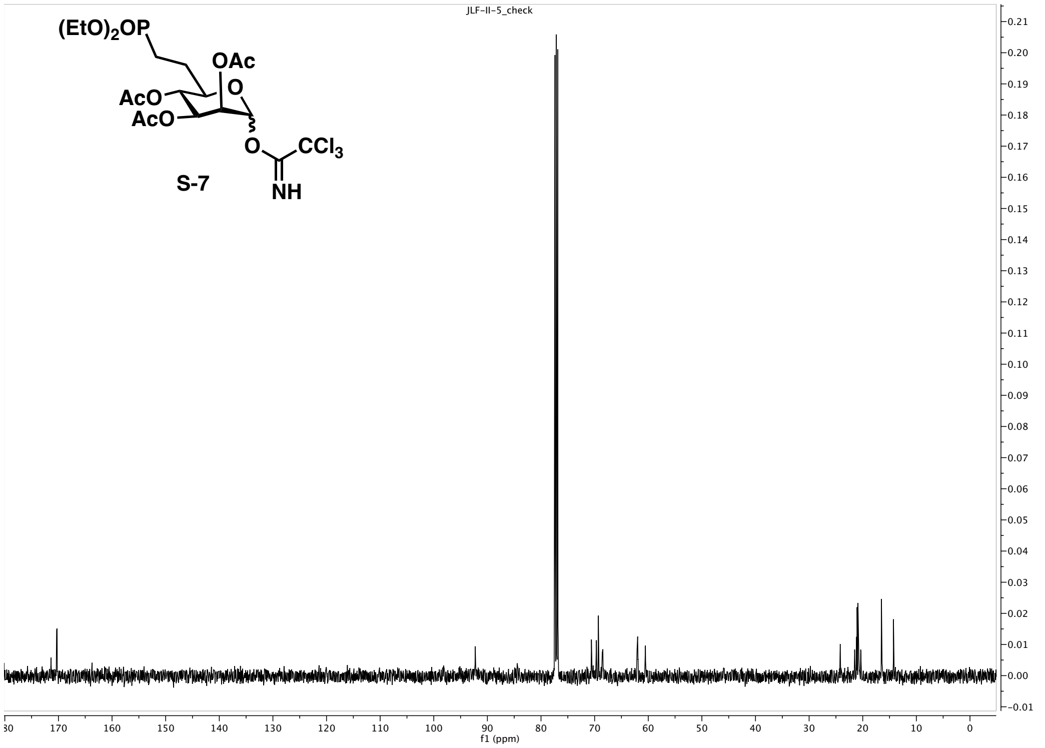


**Figure S-A.13.** $^{13}C$ NMR (100 MHz, $CDCl_3$) spectrum for compound **S-7**.

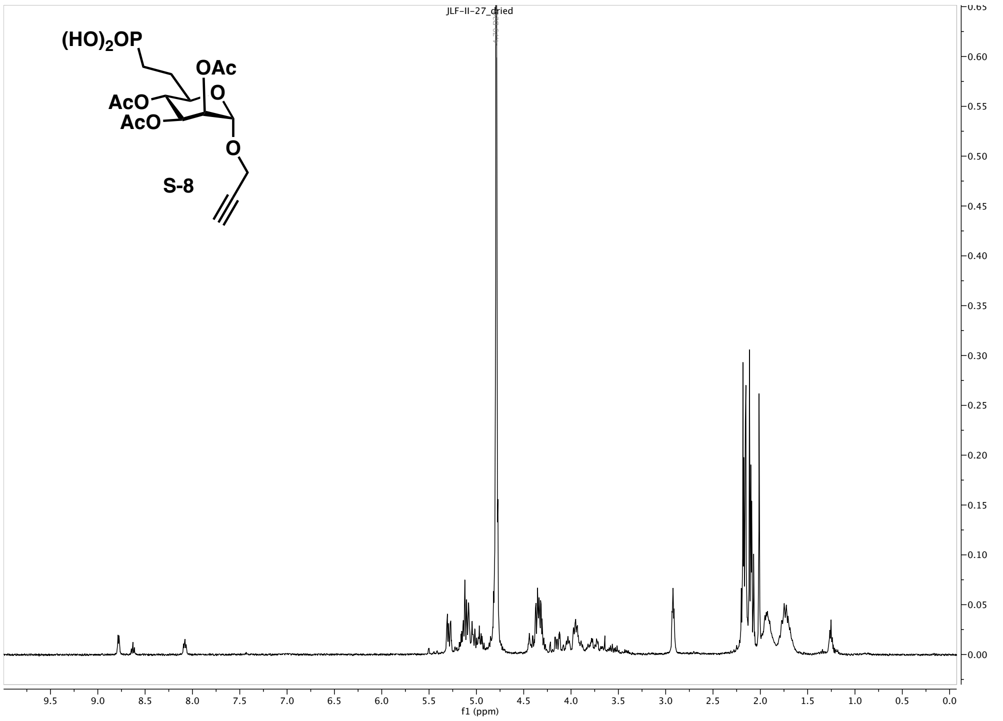


**Figure S-A.14.** $^{1}H$ NMR (500 MHz, $D_2O$) spectrum for compound **S-8**.

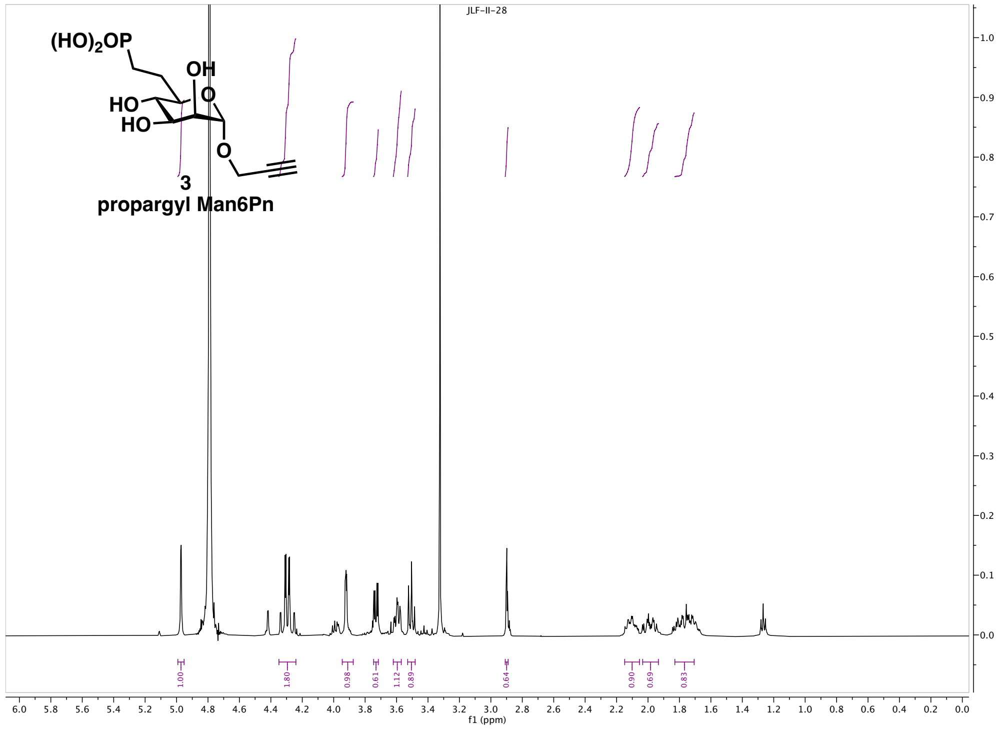


**Figure S-A.15.** $^{1}H$ NMR (500 MHz, $D_2O$) spectrum for compound **3**.

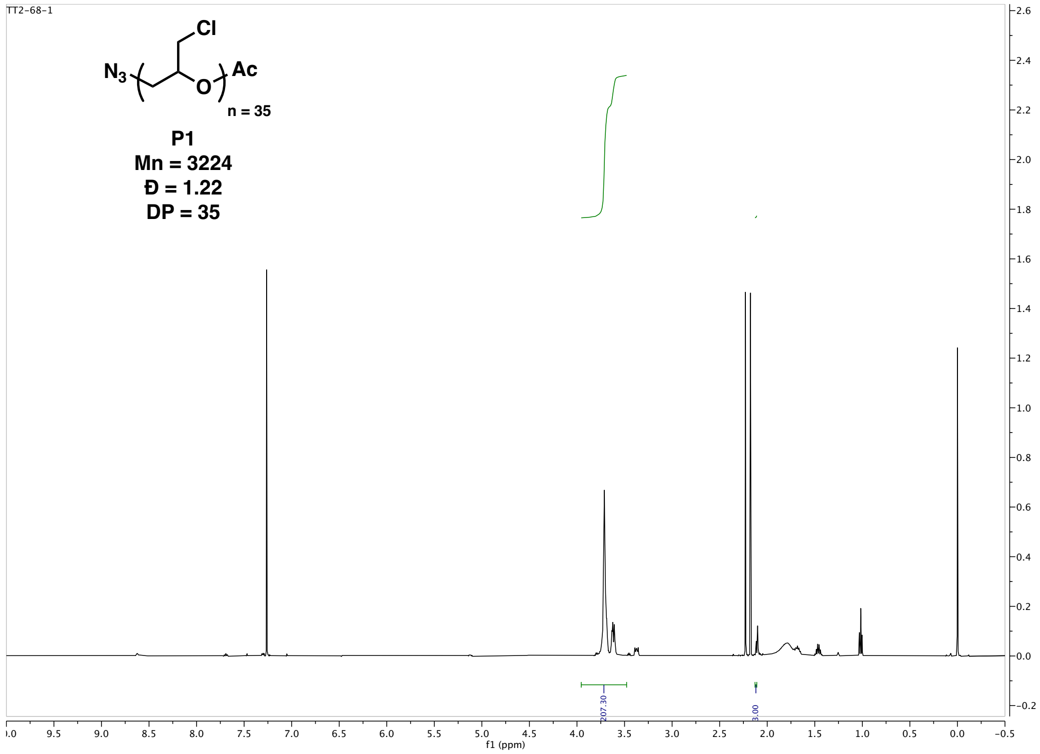


**Figure S-A.16.** $^{1}H$ NMR (500 MHz, $CDCl_3$) spectrum for polymer **P1**.

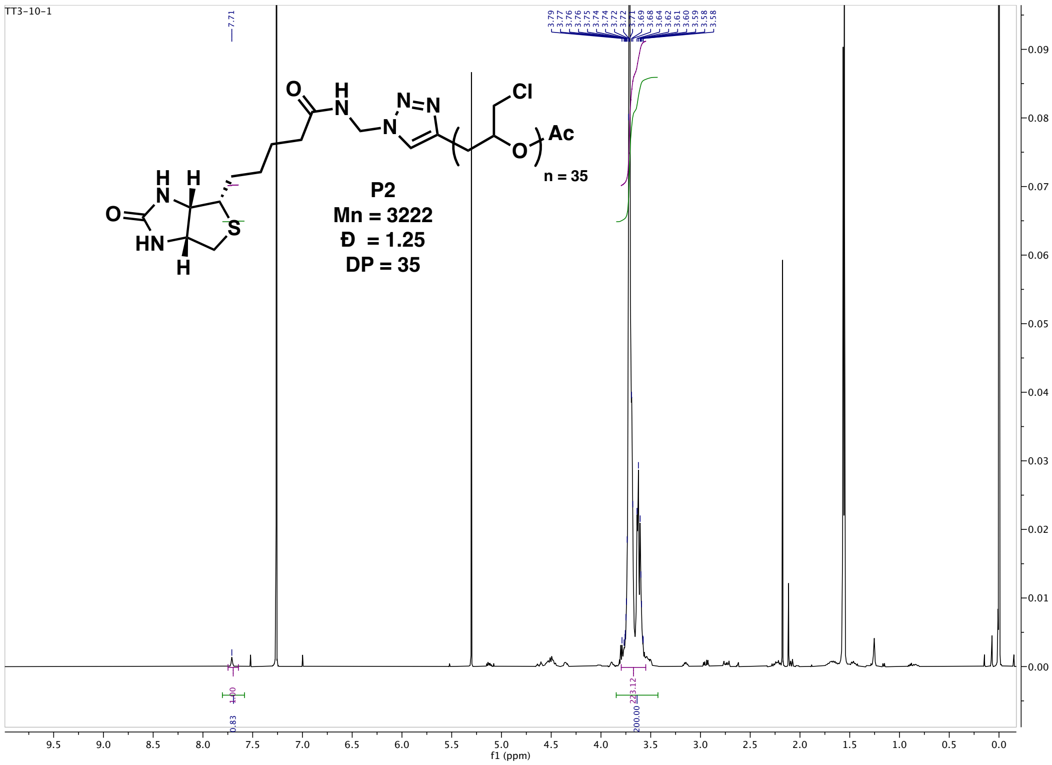


**Figure S-A.17.** $^{1}$H NMR (400 MHz, $CDCl_3$) spectrum for polymer **P2**.

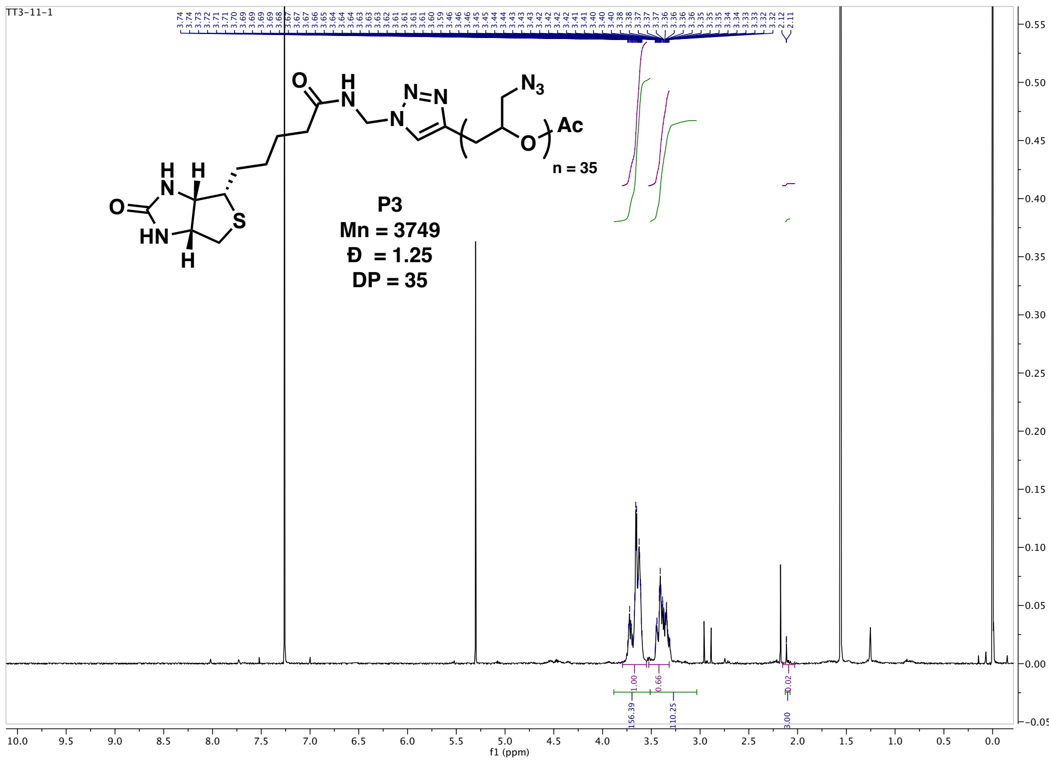


**Figure S-A.18.** $^{1}$H NMR (400 MHz, $CDCl_3$) spectrum for polymer **P3**.

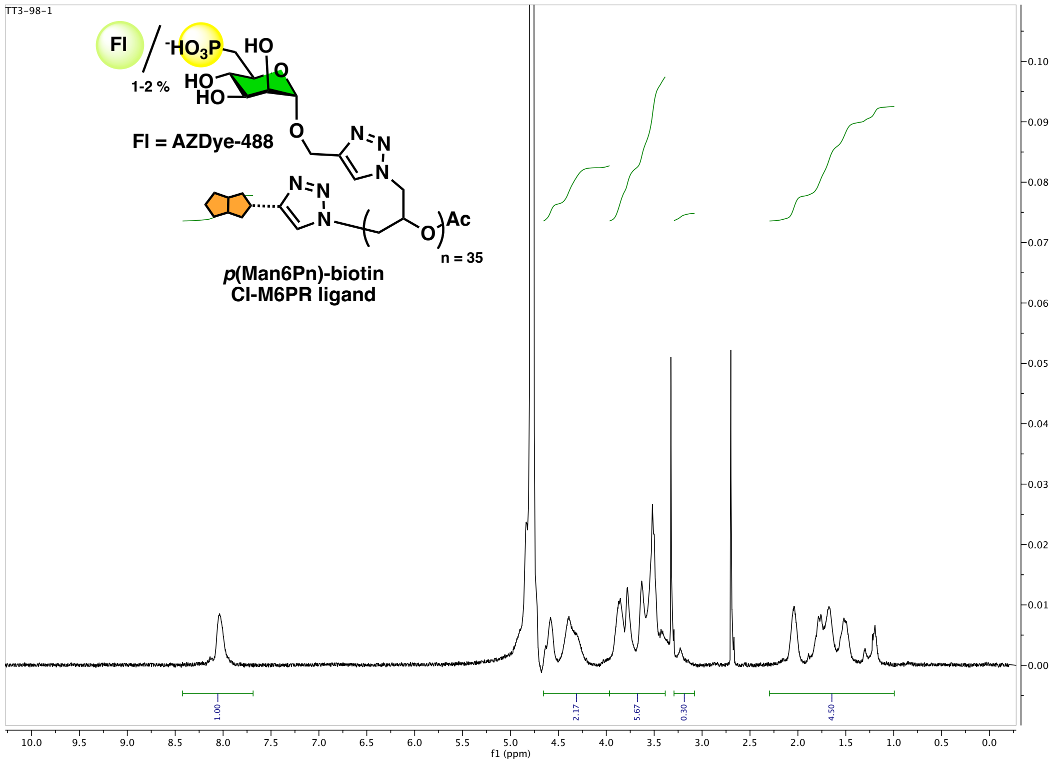


**Figure S-A.18.** $^{1}H$ NMR (400 MHz, $D_2O$) spectrum for polymer ***p*(M6Pn)-biotin**.

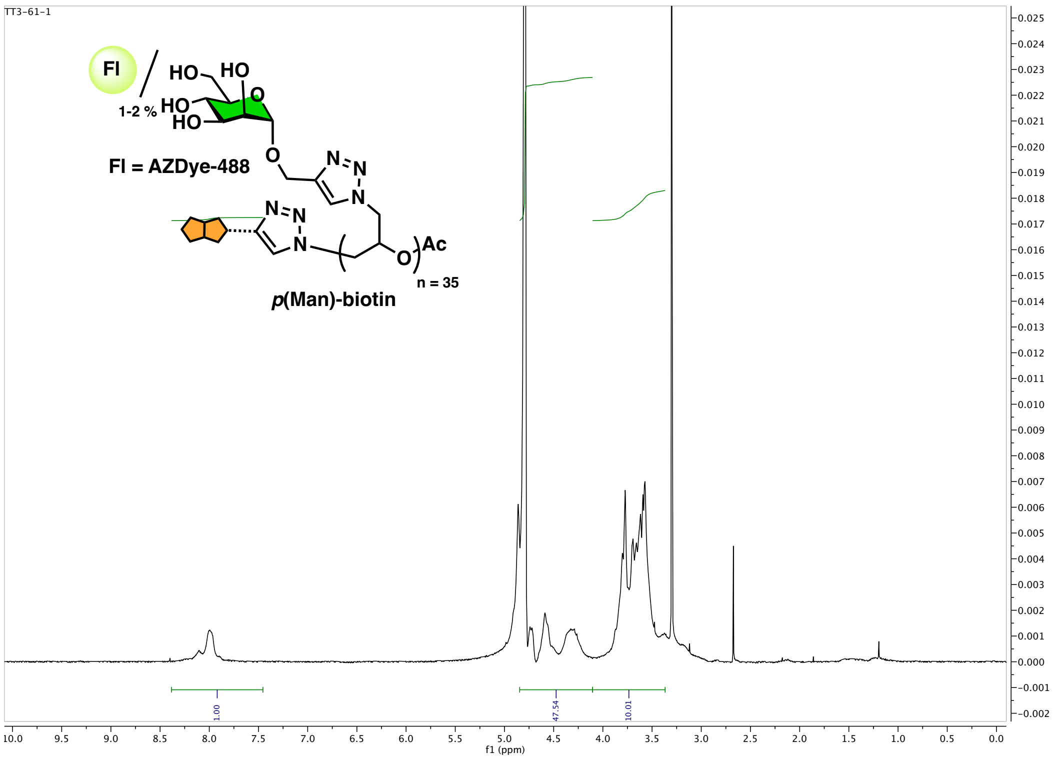


**Figure S-A.19.** $^{1}H$ NMR (400 MHz, $D_2O$) spectrum for polymer ***p*(Man)-biotin**.
